# Mitochondrial reverse electron transport in myeloid cells perpetuates neuroinflammation

**DOI:** 10.1101/2024.01.03.574059

**Authors:** L. Peruzzotti-Jametti, C.M. Willis, R. Hamel, G. Krzak, J.A. Reisz, H.A. Prag, V. Wu, Y. Xiang, A.M.R. van den Bosch, A.M. Nicaise, L. Roth, G.R. Bates, H. Huang, A.E. Vincent, C. Frezza, C. Viscomi, J.C. Marioni, A. D’Alessandro, Z. Takats, M.P. Murphy, S. Pluchino

## Abstract

Sustained smouldering, or low grade, activation of myeloid cells is a common hallmark of several chronic neurological diseases, including multiple sclerosis (MS)^1^. Distinct metabolic and mitochondrial features guide the activation and the diverse functional states of myeloid cells^2^. However, how these metabolic features act to perpetuate neuroinflammation is currently unknown. Using a multiomics approach, we identified a new molecular signature that perpetuates the activation of myeloid cells through mitochondrial complex II (CII) and I (CI) activity driving reverse electron transport (RET) and the production of reactive oxygen species (ROS). Blocking RET in pro-inflammatory myeloid cells protected the central nervous system (CNS) against neurotoxic damage and improved functional outcomes in animal disease models *in vivo*. Our data show that RET in myeloid cells is a potential new therapeutic target to foster neuroprotection in smouldering inflammatory CNS disorders^3^.

---

In MS, chronic active, slowly expanding, smouldering lesions characterised by the accumulation of myeloid cells at the lesion edge are associated with brain volume loss, irreversible disability, and disease progression^4,5^. Persistently activated myeloid cells are a continuous source of neurotoxic factors, including tumour necrosis factor-α, interleukin-1β, nitric oxide (NO), and ROS, causing remyelination failure and secondary neuronal damage^6^.

Extra- and intra-cellular metabolites, as well as mitochondrial respiratory complexes, are known to control myeloid immune responses^7,8^. Under inflammatory conditions, elevated intracellular succinate levels promote a switch from the normal forward electron transport (FET) along the respiratory chain to RET through mitochondrial complex I (CI)^7^. This mechanism, which requires a high proton motive force and a reduced coenzyme Q (CoQ) pool^9^, effectively repurposes mitochondria away from the production of adenosine triphosphate (ATP) towards the generation of superoxide that goes on to form hydrogen peroxide and other ROS, together called mitochondrial ROS (mtROS)^7^.

Inhibition of CII, via the reversible inhibitors itaconate or malonate, limits RET-induced mtROS production and promotes anti-inflammatory effects in myeloid cells *in vitro*^10,11^. Similarly, blocking CII or CI activity protects against RET-mediated mtROS damage in the infarcted heart *in vivo*^12,13^. However, the potential of RET in perpetuating the activation of myeloid cells in the context of smouldering inflammatory CNS diseases is largely unexplored.

To investigate how microglia and CNS-infiltrating myeloid cells perpetuate CNS inflammation, we used *ex vivo* single-cell RNA sequencing (scRNAseq) and liquid chromatography-mass spectrometry (LC-MS)-based analyses of *Cx3cr1*YFP^CreERT2^:R26^tdTomato^ mice^14,15^ with MOG_35-55_ experimental autoimmune encephalomyelitis (EAE), a model of chronic MS (**Fig. 1A, Extended Data Fig. 1**). RFP^+^YFP^+^ microglia and RFP^-^YFP^+^ infiltrating myeloid cells were isolated via FACS from the spinal cord of acute (A)-EAE (3 days after disease onset) and chronic (C)-EAE (50 days post immunisation, dpi) mice. Healthy (non-immunised) *Cx3cr1*YFP^CreERT2^:R26^tdTomato^ mice were used as controls (Ctrl).

**Fig. 1.**
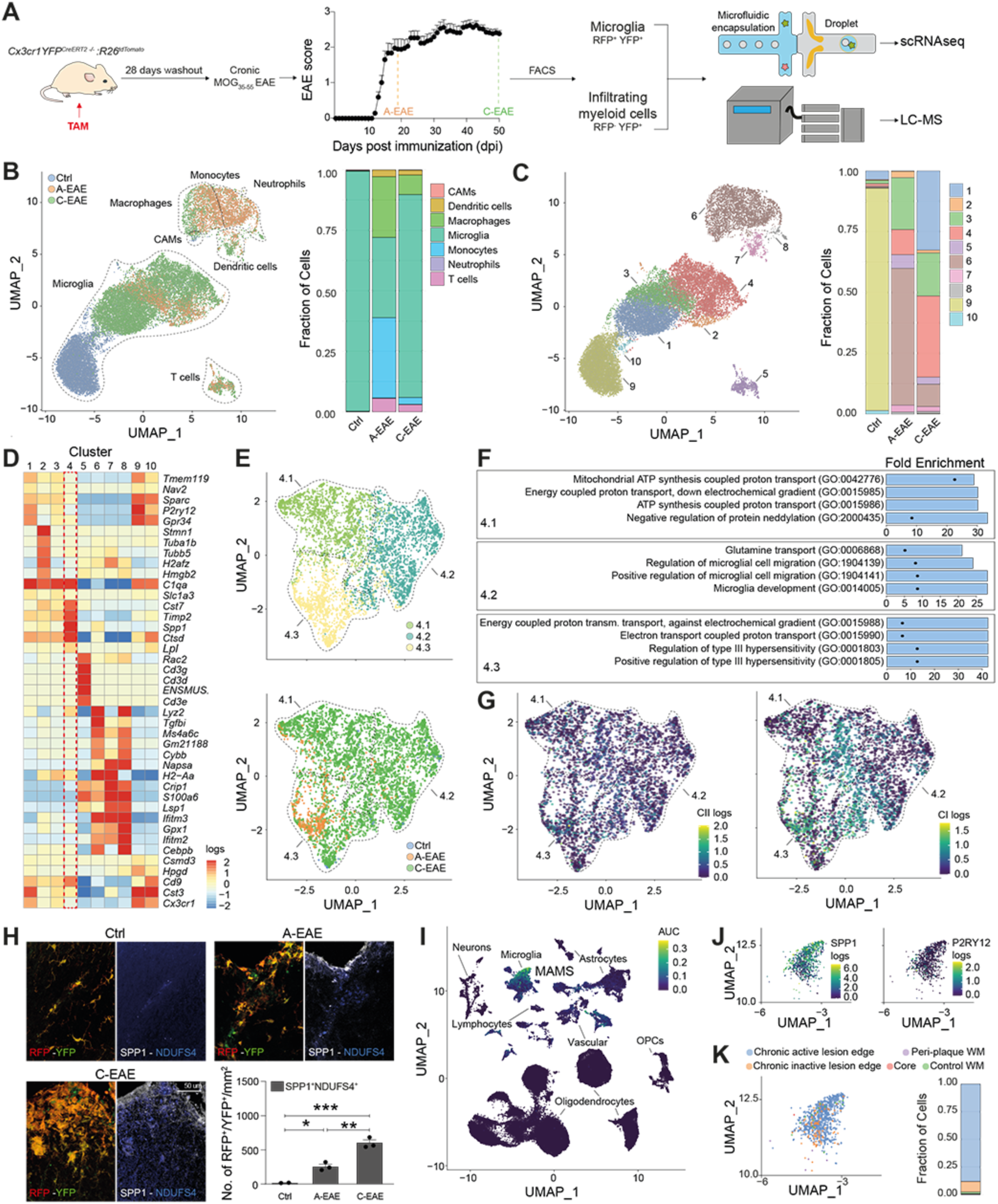
Microglia develop increased mitochondrial CI expression during EAE. (**A**) Experimental design for the *ex vivo* scRNAseq and LC-MS studies. TAM: tamoxifen. (**B**) ScRNAseq UMAP plot obtained from 22,148 cells coloured by EAE stage (Ctrl: 6,205 cells; A-EAE: 3,648 cells; C-EAE: 12,295 cells) and corresponding fraction of cell types. CAMs: CNS-associated macrophages. (**C**) UMAP plot coloured by clusters and corresponding fraction of cells in each EAE stage. (**D**) Grouped heatmap of the top differentially expressed genes for the identified clusters. Dotted red boxes highlight Cluster 4 DAM. ENSMUS: ENSMUSG00000076498. (**E**) UMAP plots of the unsupervised subcluster analysis of Cluster 4 DAM coloured by subcluster (top) and EAE stage (bottom). (**F**) Top 4 GO terms (by fold enrichment) of Cluster 4 DAM subclusters. (**G**) UMAP plots of Cluster 4 DAM subclusters coloured by the scaled median counts of mitochondrial CII (left) and CI (right). (**H**) Representative confocal imaging and quantification of EAE lesions showing the number of SPP1^+^ Cluster 4 DAM expressing the NADH ubiquinone oxidoreductase iron-sulfur protein 4 (NDUFS4). *P<0.05, **P<0.01, ***P<0.001 (one-way ANOVA). (**I**) The transcriptional profile of Cluster 4 DAM was consistent with a population of microglia (MAMS) found in progressive MS patients^18^. UMAP coloured by the area under the curve (AUC) scoring of similarity. OPCs: oligodendrocyte progenitor cells. (**J)** UMAP showing the expression of *SPP1* and of the homeostatic gene *P2RY12* in MAMS. **(K**) UMAP and bar chart showing the localisation of MAMS in progressive MS lesions and controls.

ScRNAseq revealed a prevalence of infiltrating myeloid cells in A-EAE, while C-EAE showed a prevalence of microglia (**Fig. 1B, Extended Data Fig. 1, Supplementary Data 1**). Unsupervised clustering analysis of the integrated dataset identified 10 cell clusters between Ctrl, A-EAE, and C-EAE (**Fig. 1C**). We found that the relative proportion of cells with a transcriptional signature reminiscent of microglia returning to homeostasis (hMG; Cluster 1) increased from 0.2% in A-EAE to 33.3% in C-EAE (**Fig. 1C**). Cluster 4, which identifies disease associated microglia (DAM), also showed a ∼3.2-fold increase in C-EAE (33.4%) *vs* A-EAE (10.3%) and became the dominant cluster (∼40% of total microglia) in C-EAE (**Fig. 1C**).

Cluster 4 DAM was characterised by the increased expression of the DAM genes *Spp1* (**Extended Data Fig. 2)***, Cst7, Timp2,* and *Ctsd* (**Fig. 1D**)^16^. Tissue pathology of spinal cord inflammatory lesions confirmed the presence of SPP1^+^ DAM, which were predominantly comprised of microglia and significantly increased in C-EAE (**Extended Data Fig. 2**). Further transcriptomic analyses showed that Cluster 4 DAM were also characterised by differentially expressed genes (DEGs) related to glycolysis *(e.g., Aldoa*, *Gapdh*) and oxidative phosphorylation *(e.g., Uqcrh*, *Cox6a2*) (**Supplementary Data 1, Extended Data Fig. 3**).

Subclustering analysis of Cluster 4 DAM identified three subclusters (**Fig. 1E**). Subcluster 4.1 DAM were defined by genes involved in iron metabolism (*Fth1, Flt1*) and glycolysis (*Aldoa*, *Gapdh*) (**Supplementary Data 2**). Subcluster 4.2 DAM were defined by genes associated with microglial migration (*Cx3cr1*) and identity (*Zeb2*) (**Supplementary Data 2**). Subcluster 4.3 DAM were defined by higher expression of genes of CI (*Nd4, Nd2*), complex IV (*COX1-3*) and complex V (*ATP6*) (**Supplementary Data 2**). GO-term analysis of Subclusters 4.1 and 4.3 DAM showed enrichment in pathways associated with energy coupled proton transport, both generating and utilising the proton electrochemical potential gradient, respectively (**Fig. 1F, Supplementary Data 2**).

Given the functional role of CII and CI in proton transport and ROS generation in pro-inflammatory myeloid cells^9^, genes encoding for these two mitochondrial complexes were further analysed. While we found no changes in the expression of genes encoding the CII subunits during EAE, in Cluster 4 (**Extended Data Fig. 3**), or in Subclusters 4.1-3 (**Fig 1G**), we discovered instead an upregulation of genes encoding the CI subunits in A-EAE, both in Cluster 4 DAM (**Extended Data Fig. 3)** and in Cluster 6 infiltrating myeloid cells (**Extended Data Fig. 3** and 4). CI subunits expression was further increased in Cluster 4 DAM (**Extended Data Fig. 3**) and Subclusters 4.1-3 isolated from C-EAE mice (**Fig. 1E,G**). *In situ* verification confirmed a 7.6-fold increase of the number of SPP1^+^ DAM expressing the NADH-ubiquinone oxidoreductase subunit of CI, NDUFS4^17^, (SPP1^+^/NDUFS4^+^) in the spinal cord of A-EAE mice *vs* Ctrl, and a further 2.4-fold increase of SPP1^+^/NDUFS4^+^ DAM in C-EAE mice *vs* A-EAE (**Fig. 1H**).

Re-analysis of publicly available single nuclei RNA sequencing data^18^ allowed the identification of a cluster of human microglia activated in progressive MS (MAMS), which displayed a transcriptional profile reminiscent of our own mouse Cluster 4 DAM (**Fig. 1I, Extended Data Fig. 5**). Compared to the other human microglial clusters, MAMS were characterised by the expression of *SPP1* (**Fig. 1J**), *APOE*, *MAFB*, and the CI gene (*MT-ND3*) (**Supplementary Data 2**), while expressing low levels of the homeostatic gene *P2RY12* (**Fig. 1J**). MAMS were nearly absent in the white matter of control patients, whereas they were mostly restricted (88% of the total MAMS) to the edge of slowly expanding, smouldering, chronic active lesions (CAL) of progressive MS, comprising 28% of all CAL microglia (**Fig. 1K**). These data suggest a key role for MAMS in the pathobiology of smouldering lesions and MS disease progression.

So far, our work identified a cluster of persistently activated DAM with high expression of CI genes and proteins, which increases during chronic smouldering CNS inflammation in an animal disease model, and is found almost exclusively at the edge of smouldering CAL of people with progressive MS.

To gain insights onto the metabolic features of microglia and infiltrating myeloid cells that act to perpetuate chronic CNS inflammation, we performed LC-MS analysis of the intracellular metabolome of *ex vivo* isolated myeloid cells. We found a clear separation based on the partial least squares-discriminant analysis and a differential abundance of intracellular metabolites based on the cell type and stage of EAE (**Extended Data Fig. 6, Supplementary Data 3**).

A-EAE microglia showed increased intracellular levels of itaconate, phosphocreatine (an ATP buffer)^19^, the ROS scavengers ascorbate and dehydroascorbate, and glutathione disulfide (GSSG), which arises as a consequence of antioxidant reactions^20^ *vs* Ctrl (**Fig. 2A-B, Supplementary Data 3**). Laser desorption - rapid evaporative ionisation mass spectrometry (LD-REIMS) of spinal cord sections confirmed the higher abundance of itaconate and ascorbate within white matter inflammatory infiltrates *in situ* (**Extended Data Fig. 7, Supplementary Data 4**). Correlation analysis of itaconate within the entire LC-MS dataset showed a direct correlation between itaconate levels and the ROS scavengers ascorbate and dehydroascorbate (**Extended Data Fig. 7**). Instead, C-EAE microglia showed significantly lower amounts of GSSG, phosphocreatine, ATP (*vs* A-EAE), and lower itaconate, but significantly increased intracellular levels of creatine and L-citrulline (**Fig. 2C-E, Supplementary Data 3**). These findings are suggestive of elevated NO, peroxynitrite, and ROS generation in C-EAE microglia^21^.

**Fig. 2.**
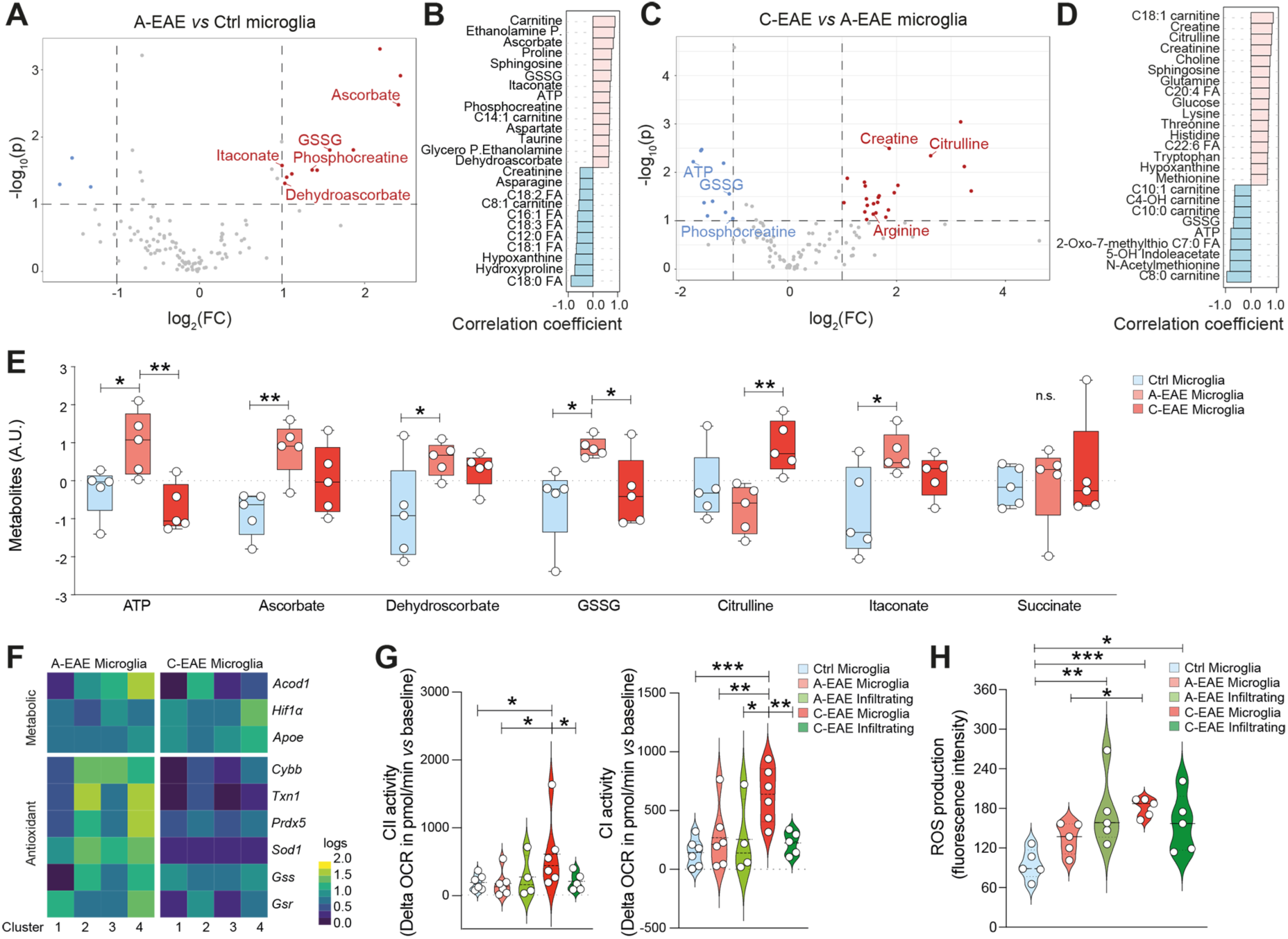
Metabolomic analysis of myeloid cells show changes in energy metabolism and oxidative stress in EAE. (**A**) Volcano plot of the metabolites significantly altered in A-EAE microglia *vs* Ctrl. (**B**) Corresponding correlation analysis of metabolites indicative of A-EAE microglia *vs* Ctrl. (**C**) Volcano plot of the metabolites significantly altered in C-EAE *vs* A-EAE microglia. (**D**) Corresponding correlation analysis of metabolites indicative of C-EAE *vs* A-EAE microglia. (**E**) Bar plots of selected relevant metabolites. P.: phosphate, FA: fatty acids. A.U. arbitrary units. (**F**) Heatmap showing genes (from our scRNAseq dataset) involved in itaconate synthesis (*Acod1*), glycolytic switch (*Hif1α*), DAM phenotype (*Apoe*), glutathione (*Gss*, *Gsr*), and antioxidant response in the microglial clusters isolated from A-EAE *vs* C-EAE mice. (**G**) CII and CI activity in *ex vivo* FACS isolated microglia and infiltrating myeloid cells from EAE mice. OCR: Oxygen consumption rate. (**H**) Quantification of fluorescence intensity of the CellROX probe signal in FACS isolated microglia and infiltrating myeloid cells from EAE mice. *P<0.05, **P<0.01, ***P<0.001 (one-way ANOVA).

To link these metabolic changes with the expression of relevant genes from our scRNAseq dataset, we focused on the clusters of microglia isolated from A-EAE and C-EAE mice (**Fig. 2F**). We found that the expression of *Aconitate decarboxylase 1* (*Acod1*), which codes for the enzyme that synthesises itaconate^11^, was increased in Cluster 4 DAM in A-EAE but decreased in C-EAE mice. Conversely, *Hif1α*, which is involved in the switch to aerobic glycolysis in myeloid cells^22^, and the DAM marker *Apoe,* were both increased in Cluster 4 DAM during C-EAE. Genes known to control the antioxidant response to ROS (e.g., *Cybb, Txn1, Prdx5, Sod1*) and glutathione synthesis/reduction in microglia (*Gss, Gsr*)^23,24^ were also increased in Cluster 4 DAM in A-EAE but decreased in C-EAE.

To further connect these metabolic and transcriptional features with the function of CII and CI, we applied *ex vivo* metabolic flux analysis to both microglia and infiltrating myeloid cells at different stages of disease. We found that C-EAE microglia had significantly higher levels of both CII and CI activity (**Fig. 2G**), as well as higher levels of oxidative stress (**Fig. 2H**), suggesting elevated mtROS production^7^ in these cells during C-EAE.

Based on these integrated data, we propose that after transitioning from A-EAE to C-EAE, microglia display increased CII-CI activity and oxidative stress, coupled with lower itaconate levels and ATP, which support a repurposing of their mitochondria towards ROS generation.

To investigate whether CII and CI act via RET to determine and amplify the increased oxidative stress seen in chronic smouldering CNS inflammation, we then developed an *in vitro* model of RET induction (RET^+^) in pro-inflammatory microglia. RET was induced by treating LPS/IFNγ stimulated microglial cells with oligomycin, which blocks complex V and increases the mitochondrial proton motive force, and succinate, to provide a substrate for oxidation by CII, reduce the CoQ pool, and sustain high mitochondrial proton motive force (**Fig. 3A**). RET induction amplified the production of mtROS in pro-inflammatory microglia (*vs* RET^-^) (**Fig. 3B,C**). Treatment of RET^+^ pro-inflammatory microglia with the CI inhibitor rotenone significantly hampered mtROS production and prevented the increase of mitochondrial membrane potential (**Fig. 3B-D**). These data confirm a central role for CI during RET *in vitro*, which acts to amplify mtROS production in pro-inflammatory microglia.

**Fig. 3.**
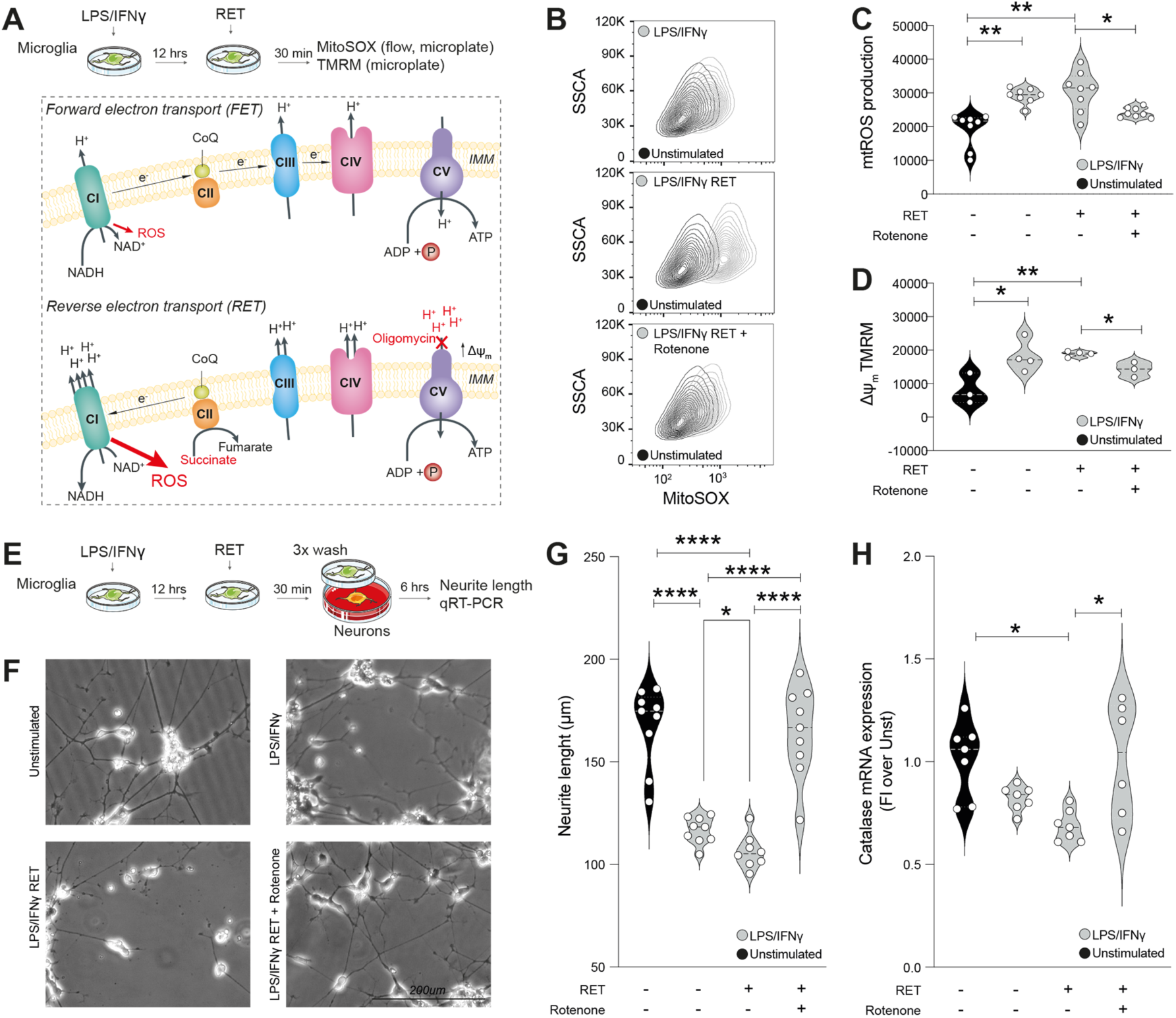
Blocking RET in microglia reduces excessive mtROS production and neurotoxicity *in vitro*. (**A**) Experimental design for the *in vitro* RET studies on LPS/IFNγ stimulated microglia. IMM: inner mitochondrial membrane. (**B**) Qualitative flow cytometry of mtROS production assessed via the MitoSOX probe in unstimulated and LPS/IFNγ stimulated microglia after RET induction. Rotenone is given to block CI activity. (**C-D**) Quantification of mtROS production via the MitoSOX probe (C) and mitochondrial membrane potential via the TMRM probe (D) in LPS/IFNγ stimulated microglia after RET induction. (**E**) Experimental design for the *in vitro* microglia-neuronal trans well co-cultures. (**F-G**) Representative images (F) and quantification of neurite length (G) after co-culture with LPS/IFNγ stimulated microglia after RET induction. (**F**) Catalase mRNA expression in neuronal cells co-cultured with LPS/IFNγ stimulated microglia after RET induction. *P<0.05, **P<0.01, ***P<0.001 (one-way ANOVA).

To test the pathogenic role of RET^+^ pro-inflammatory microglia we then co-cultured them with SH-SY5Y neurons using a trans well co-culture system that avoids cell-to-cell contacts (**Fig. 3E**). We observed a significant loss of neurites in SH-SY5Y neurons co-cultured with RET^-^ pro-inflammatory microglia, which was further amplified by RET^+^ pro-inflammatory microglia (**Fig. 3F-G**). We also found a significant downregulation in *CAT* (catalase) mRNA expression levels in SH-SY5Y neurons caused by RET^+^ pro-inflammatory microglia, which is supportive of ROS mediated damage via phosphatidylinositol 3 kinase/Akt signalling^25^ (**Fig. 3H**). Blocking CI activity in RET^+^ pro-inflammatory microglia entirely rescued neurite length and *CAT* expression in SH-SY5Y neurons (**Fig. 3G-H**). Altogether, these data suggest that blocking RET in pro-inflammatory microglia via CI inhibition protects from excessive mtROS-associated neurotoxicity *in vitro*.

To confirm the relevance of these findings in CNS inflammation *in vivo*, we induced MOG_35-55_ EAE in *Nd6* mice, which carry a point mutation in the mitochondrial gene *Nd6* that blocks RET while conserving normal FET^26^. *Nd6* mice showed a significantly lower disease severity throughout the entire course of EAE *vs* wild type (WT) mice (**Fig. 4A**). *Ex vivo* scRNAseq of the entire CNS of *Nd6* EAE mice at 30 dpi (an intermediate time point between A- and C-EAE) revealed 14 cell clusters (**Fig. 4B** and **Extended Data Fig. 8**). Clusters 4, 10 and 11 were classified as DAM, having low expression of hMG genes (*Sparc, P2ry12, Cd33, Tmem119*), and high expression of *Spp1* and other DAM genes (i.e., *Apoe, Tyrobp, Ctsd,* as well as *Trem2, Lpl, Clec7a*)^16^ (**Fig. 4C**). These DAM clusters were also characterised by GO terms related to microglial activation and cytotoxicity (**Supplementary Data 5**). Importantly, we found that *Nd6* EAE mice had a ∼1.8-fold and ∼2.6-fold decrease of Cluster 10 and 11 DAM *vs* WT (6.8 *vs* 12.9% and 6.1 *vs* 16.2%, respectively) (**Fig. 4B**), and this was associated with a significant reduction of *in vivo* mtROS production^27^ (**Fig. 4D**).

**Fig. 4.**
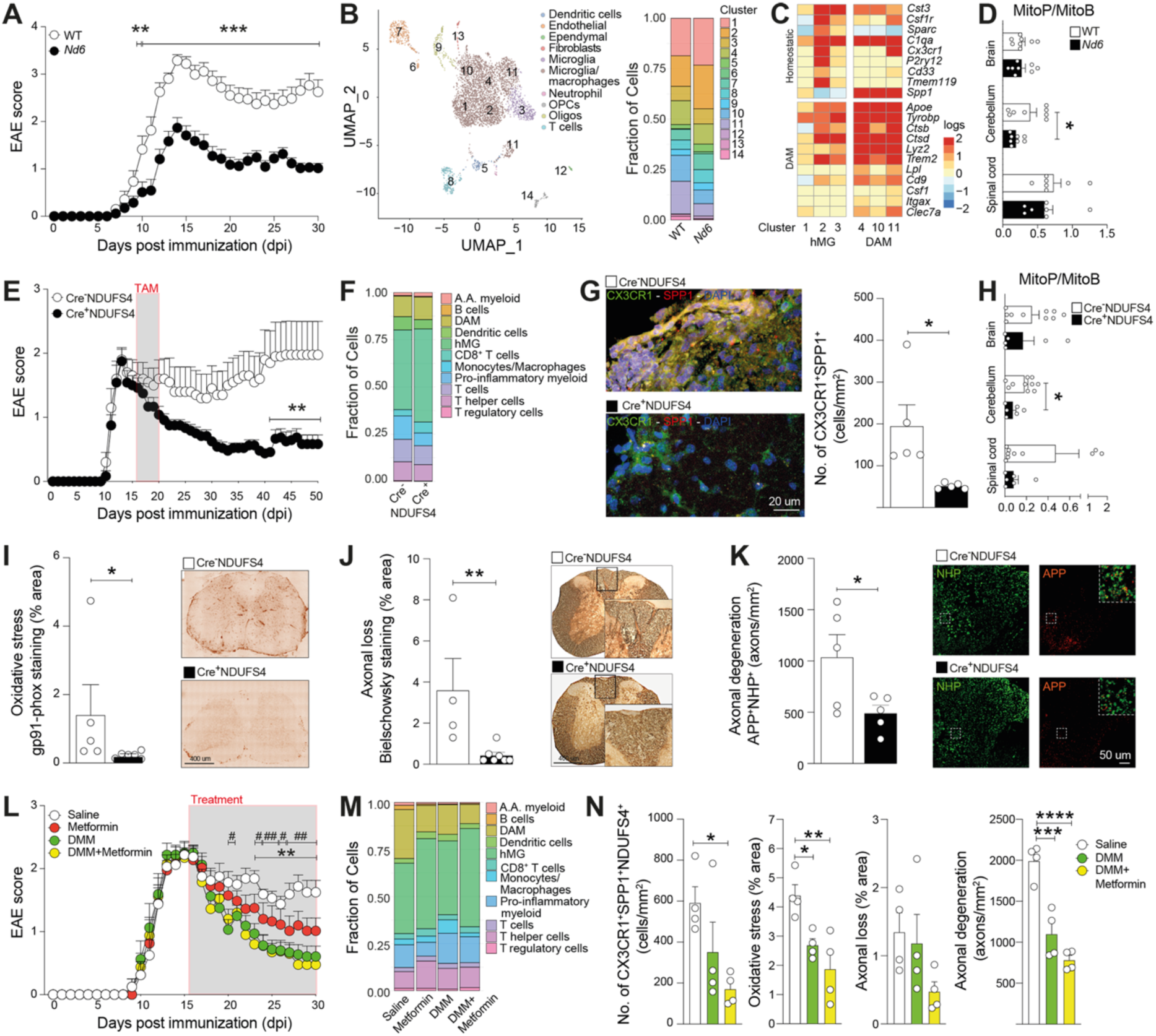
Blocking RET *in vivo* ameliorates EAE and reduces secondary axonal damage. (**A**) EAE scores of WT and *Nd6* mice at 30 dpi. **P<0.01, ***P<0.001 (two-way ANOVA). (**B**) ScRNAseq UMAP plot coloured by cell type with superimposed clusters (left), and corresponding fraction of cells (right). OPCs: oligodendrocyte progenitor cells. (**C**) Heatmap of selected genes in hMG and DAM clusters. (**D**) Quantification of *in vivo* mtROS production via MitoP/MitoB. *P≤0.05 (unpaired t-test). (**E**) EAE scores of Cre^-^NDUFS4 and Cre^+^NDUFS4 mice. **P<0.01 (two-way ANOVA). (**F**) CyTOF analysis of immune cell types at 50 dpi. A.A.: alternatively activated (**G**) Confocal imaging and quantification of CX3CR1^+^SPP1^+^ cells in EAE lesions at 50 dpi. *P≤0.05 (unpaired t-test). (**H-K**) *In vivo* mtROS production (H), oxidative stress (I), axonal loss (J), and axonal degeneration (K, insets are merged images, NHP: neurofilament heavy polypeptide, APP: amyloid precursor protein) at 50 dpi. *P≤0.05, **P<0.01 (unpaired t-test). (**L**) EAE scores of mice treated with Metformin, DMM, DMM+Metformin *vs* saline controls. #P<0.05, ##P<0.01 DMM *vs* saline; **P<0.01 DMM+Metformin *vs* saline (two-way ANOVA). (**M**) CyTOF analysis of immune cell types at 30 dpi. (**N**) Quantification of CX3CR1^+^SPP1^+^NDUFS4^+^ cells, oxidative stress, axonal loss, and axonal degeneration in EAE mice at 30 dpi. *P<0.05, **P<0.01, ***P<0.001, ****P<0.0001 (one-way ANOVA).

To selectively interfere with CI activity in myeloid cells only, and to target the transition between A-EAE and C-EAE, we next generated a tamoxifen (TAM) inducible transgenic mouse line that allows for the timed knockout of *Ndufs4*^28^ in myeloid cells *in vivo* (**Extended Data Fig. 9**). We induced MOG_35-55_ EAE in *Cx3cr1*YFP^CreERT2^*:Ndufs4*^flox/flox^ mice (Cre^+/-^ NDUFS4) and administered TAM one week after the onset of EAE symptoms.

We found that Cre^+^NDUFS4 EAE mice had significantly lower disease severity at the C-EAE stage *vs* Cre^-^NDUFS4 mice (**Fig. 4E**). *Ex vivo* mass cytometry analysis of CD45^+^ spinal cord cells (**Supplementary Data 6**) revealed a 16.9% increase in the number of hMG in Cre^+^NDUFS4 mice (**Fig. 4F**), while DAM clusters displayed significantly lower CII and CI expression *vs* Cre^-^NDUFS4 mice (**Extended Data Fig. 9**). Tissue pathology of spinal cord inflammatory lesions showed that Cre^+^NDUFS4 mice had a significant reduction of CX3CR1^+^SPP1^+^ DAM (**Fig. 4G**), coupled with lower levels of *in vivo* mtROS (**Fig. 4H**), tissue oxidative stress^24^ (**Fig. 4I**), and axonal loss and degeneration^29^ *vs* Cre^-^NDUFS4 mice (**Fig. 4J-K**).

We next performed an *in vitro* drug testing and *in vivo* pharmacokinetic study of selected CII and CI inhibitors (**Extended Data Fig. 10**). We then induced MOG_35-55_ EAE in C57/Bl6 mice and applied daily intraperitoneal (IP) injections of either the CII inhibitor dimethyl malonate (DMM), or Metformin (which is known to reduce the generation of ROS at CI)^30^, or the combination of DMM+Metformin (**Fig. 4L**). Mice were treated starting one week after disease onset to target the transition between A-EAE and C-EAE. We found that DMM and DMM+Metformin induced the most significant recovery of clinical signs at 30 dpi. *Ex vivo* mass cytometry analysis of CD45^+^ spinal cord cells showed a striking effect of DMM+Metformin on the frequency of hMG (1.4 and 1.3-fold increase *vs* Ctrl and DMM, respectively) and DAM (2.5 and 1.6-fold decrease *vs* Ctrl and DMM, respectively) (**Fig. 4M**). DMM+Metformin also led to a significant reduction of CII and CI expression in DAM clusters (**Extended Data Fig. 10**). Finally, tissue pathology confirmed that DMM+Metformin led to a significant reduction of SPP1^+^NDUFS4^+^ DAM, which was coupled with lower levels of tissue oxidative stress, axonal loss and degeneration in the spinal cord (**Fig. 4N**).

In conclusion, our data suggest that blocking RET in myeloid cells using both genome engineering and small molecules targeting mitochondrial CII and CI function is a promising avenue to interfere with the irreversible accumulation of secondary neurological damage caused by excess mtROS in smouldering CNS inflammation.

## Methods

### Mice and tamoxifen treatment

The *Cx3cr1*YFP^CreERT2^:R26^tdTomato^ fate mapping mouse was generated by crossing B6.129P2(Cg)-Cx3cr1tm2.1(cre/ERT2)Litt/WganJ^31^ with B6.Cg-Gt(ROSA)26Sortm9(CAG-tdTomato)Hze/J), as previously described^14^. To induce RFP expression, tamoxifen (TAM) (Sigma-Aldrich) was diluted in corn oil (Sigma-Aldrich) at a concentration of 25 mg/ml and administered daily via intraperitoneal (IP) injections for 5 days at 0.125 mg tamoxifen/g body weight. After a washout period of 28 days to allow peripheral myeloid cells to be replaced *de novo* (and thus lose RFP expression)^32^, EAE was induced. ND6-P25L mice (here called *Nd6* mice) were generated as described^26^ and obtained from Professor Douglas Wallace (University of Pennsylvania, USA). The *Cx3cr1*YFP^CreERT2^:*Ndufs4*^flox/flox^ mice were generated by crossing B6.129P2(Cg)-Cx3cr1tm2.1(cre/ERT2)Litt/WganJ^31^ with mice with conditional alleles of the *Ndufs4* gene (exon 2 flanked by loxP sites)^28^. To knockout *Ndufs4* expression in CX3CR1^+^ cells, TAM was diluted in corn oil (as above) and administered via daily IP injections for 5 days (as above) starting one week after EAE onset. WT C57BL/6 female mice were purchased from Charles River and used to test small molecules in EAE as further described below.

### EAE induction and behaviour

C57BL/6 female mice were immunised with myelin oligodendrocyte glycoprotein (MOG) 35-55, as previously described^8^. Additional information is provided in the Supplementary Methods.

### Single-cell RNA sequencing (scRNAseq)

Mice were deeply anaesthetised with IP injections of ketamine (10 mg/ml, Boehringer Ingelheim) and xylazine (1.17 mg/ml, Bayer) and then transcardially perfused for 7 min with ice-cold artificial cerebral spinal fluid (aCSF) (NaCl 87mM, KCl 2.5mM, NaH_2_PO_4_ 1.25 mM, NaHCO_3_ 26 mM, sucrose 75 mM, glucose 20 mM, CaCl_4_ 1 mM, MgSO_4_ 7 mM), as described^33–36^. Additional information is provided in the Supplementary Methods.

### Liquid Chromatography-Mass Spectrometry (LC-MS) analysis

For *ex vivo* LC-MS, details of cell isolation are provided in the Supplementary Methods. Data analysis was performed as previously described^37,38^.

### Laser Desorption – Rapid Evaporative Ionisation MS (LD-REIMS)

Data were first pre-processed using a workflow developed in-house and described in more detail elsewhere^39^. Concisely, a signal-to-noise ratio (SNR)-based peak detection algorithm was used to reduce the three-dimensional raw data into lists of detected peaks at each pixel. After the effective data re-sampling, further reduction in data size was achieved via a semi-supervised region-of-interest (ROI) selection procedure, which performed automatic annotation of pixels to be either sample or background related according to some preliminary manual selection. This ROI mask file generated was then loaded in the nominal next step of the workflow, which aligned and mass-corrected all the detected peaks by a 2-step algorithm using user-defined references masses (m/z = 174.0408, 255.2324, 766.5392 in this case) as tissue-unrelated peaks were filtered by the custom-built SPUTNIK package. As a result, a data cube of dimension M × N was produced for each imaging run, where M is the total number of pixels and N is the length of the common mass axis, which was also shared between all runs by considering only the masses that are common in all datasets.

To probe the metabolic differences between varying regions and between images in an untargeted fashion, unsupervised colocalisation analysis based on ranked intensity correlations was performed on the pre-processed data. This effectively paired up spectral features that produced statistically significant spatial correlation, whose relative intensity distributions can be overlayed for visual inspection. All paired features extracted this way were then manually identified via literature search and online databases (HMDB)^40^, then visualised on a circle plot. To facilitate multivariate supervised analysis, further refinement of the pre-processed data was carried out by computing a weighted average of Structural Similarity Index (SSIM) and multi-SSIM for images of every spectral feature with a representative reference image; those that produced a weighted average below a perceptually defined threshold were deemed to be belonging to either background or isotopes and hence removed. The remaining data cube was then visualised by a spectral unmixing algorithm^41^ into (k=2) components (corresponding to tissue and background in image) whose abundances can be spatially mapped to give high contrast images. Spectra from certain anatomically important regions (e.g., infiltrated white matter, as seen on haematoxylin and eosin staining) were then labelled and extracted by means of manual ROI selection on the abundance image. Weighted logistic regression (LR) classification models were then built on the labelled spectra and their performance assessed using leave-one-out-cross-validation between groups. Model refinement through feature selection was then performed based on recursive feature elimination. The validity of the selected features was verified by using them to train another LR model, and then comparing the Receiver Operator Characteristics curves (ROC) of the new and original models and their respective areas under the curve (AUC). These features were considered significant in driving the separation between anatomical and pathological features within the imaging data and were identified following the same procedure as before.

### Mitochondrial membrane potential of microglial BV2 cells *in vitro*

For the *in vitro* analysis of mitochondrial membrane potential after RET induction, the tetramethylrhodamine methyl ester (perchlorate) TMRM (Cambridge Bioscience) probe was used following manufacturer’s instruction. Additional information is provided in the Supplementary Methods.

### Mitochondrial superoxide production of microglial BV2 cells *in vitro*

For the *in vitro* analysis of mitochondrial superoxide production after RET induction, the MitoSOX Red Mitochondrial Superoxide Indicator (ThermoFisher Scientific) probe was used following manufacturer’s instructions. Additional information is provided in the Supplementary Methods.

### Microglial BV2 cells and SH-SY5Y *in vitro* co-cultures

See Supplementary Methods.

### Assessment of *ex vivo* ROS production

See Supplementary Methods.

### Metabolic flux analysis

Metabolic flux measurements were obtained from myeloid cells isolated *ex vivo* from *Cx3cr1*YFP^CreERT2^:R26^tdTomato^ mice via FACS, as described for cells undergoing LC-MS analysis. Additional information is provided in the Supplementary Methods.

### Genomic PCR

The genotype of the mice was verified via genomic PCR by collecting ear notch and extracting DNA by adding DNA Releasy (15 µL, Anachem) to each sample. Additional information is provided in the Supplementary Methods.

### Small molecules for *in vivo* studies

We first ran a pharmacokinetic study to determine the optimal route of administration of dimethyl malonate (DMM) (Sigma-Aldrich) and disodium malonate (DSM) (Sigma-Aldrich) that would lead to both the lowest blood concentration and highest CNS tissue concentration. For the pharmacokinetic study, EAE mice at one week (7 days) from clinical disease onset were separated into 6 treatment groups (n=2/group): (1) IP DMM (160 mg/kg), (2) IP DSM (160 mg/kg), (3) oral (drinking water) DMM (1.5 w/v%), and (4) oral DSM (1.5 w/v%), (5) IP saline, and (6) oral saline. IP injections: mice received 5 daily IP injections of either DMM, DSM, or saline, were allowed to rest for 30 minutes, then blood was collected via the tail vein. Oral administration: mice were given 5 days of ad libitum access to DMM, DSM, or saline treated water; blood was collected via the tail vein daily for 4 days. Obtained blood was centrifuged at 10,000 × *g* for 5 minutes and the purified plasma was collected and stored at -80°C until analysis. At the end of the treatment period, the brain and spinal cord was extracted from the mice and flash frozen in liquid nitrogen (LN2). The frozen tissue was then stored at -80°C until analysis.

Malonate from blood and tissue was quantified as previously described^42^. Briefly, 20-30 mg of tissue was homogenised in 25 µL/mg extraction buffer (50% methanol, 30% acetonitrile and 20% water) and 1 nmol ^13^C_3_-malonate as an internal standard, using pre-filled bead mill tubes (ThermoFisher Scientific) in a Precellys 24 homogeniser (Bertin Instruments) (6,500 rpm; 15 s × 2, cooling on ice in between). 10 µL blood were added to 450 µL extraction buffer containing the internal standard in a microcentrifuge tube and vortexed. All samples were subsequently incubated at -20 °C for 1 hour before centrifugation at 17, 000 × g for 10 min at 4 °C. The supernatant was transferred to a microcentrifuge tube and centrifuged at the same conditions. The resulting supernatant was transferred to pre-cooled mass spectrometry vials and stored at -80 °C until analysis by LC-MS/MS.

Metformin powder (Sigma-Aldrich) was dissolved in sterile saline and given at 100mg/kg. DMM solution was mixed with saline and given at 160mg/kg. For the combination of DMM+Metformin, Metformin powder was dissolved in sterile saline and DMM was mixed with sterile saline and then combined 1:1. At one week from the onset of clinical signs of EAE, mice were randomised into four treatment groups: (1) saline, (2) metformin, (3) DMM, (4) DMM+Metformin. Mice were given daily IP injections with the treatment or control substances until the end of the study (30 dpi), when mice were further randomised for *ex vivo* tissue pathology or suspension mass cytometry (CyTOF).

### Tissue pathology

See Supplementary Methods.

### Suspension Mass Cytometry (CyTOF)

Details of tissue processing for CyTOF, up to fcs files generation, are provided in the Supplementary Methods. Fcs files were loaded using *flowCore (2.6.0).* For the small molecules EAE samples, we used the package *GdClean* (https://github.com/JunweiLiu0208/GdClean), to estimate and remove gadolinium contamination from sample. Flow sets were then converted to single cell experiment objects using *CATALYST (1.18.1)*. Using *CATALYST,* cells were assigned to one of 100 grid points on a self-organizing map and metaclustered into 20 groups via the consensus clustering method considering cell surface features only^43^. UMAP dimensionality reduction was performed using the *CATALYST* wrapper around *scater runUMAP* with the default parameters and data from cell surface features only. Each of the 100 clusters were assigned cell types based on expert knowledge of cell type-specific protein expression patterns.

### Statistical analyses

Unless otherwise stated, statistical analyses were performed with Graph Pad Prism 9 for macOS (GraphPad Software). EAE scores were compared using two-way ANOVA followed by Bonferroni multiple comparisons test. Differences between two groups were analysed using an unpaired t-test (unless otherwise stated). Differences among >2 groups were analysed using one-way ANOVA followed by Tukey’s multiple comparisons test (unless otherwise stated). Data is shown in the text and figures as mean values ± SEM (unless otherwise stated), and a p value < 0.05 was accepted as significant in all analyses (unless otherwise stated). Methods of remaining statistical analyses have been described in the relevant sections reported above.

### Ethics Statement and Data Availability

Animal research has been regulated under the Animals (Scientific Procedures) Act 1986 Amendment Regulations 2012 following ethical review by the University of Cambridge Animal Welfare and Ethical Review Body (AWERB). Animal work was covered by the PPL 80/2457 and PPL PP2135981 (to SP). Complete R analysis workflow for scRNAseq and CyTOF, including scripts for generating the figures, are available on GitHub (regan-hamel/ComplexI). ScRNAseq and CyTOF data are available at https://data.mendeley.com/v1/datasets/w5wtx43528/draft?a=d3079d50-32aa-43d4-b719-c196d8af220e. Metabolomics data are reported as Supplementary Data.

## Acknowledgments

The authors wish to acknowledge P. Chinnery, F. Dazzi, D. Franciotta, G. Griffiths, J. Smith, A. Tolkovsky, and J. Van den Ameele for their critical insights throughout this study. The authors also thank A. Tolkovsky, M. Whitehead, and D. Wallace for providing the microglial BV2 cells, neuronal SH-SY5Y cells, and the ND6 mice, respectively. We also thank H. Bridges for her technical assistance with the setup and analysis of the *ex vivo* metabolic flux experiments, D. Trajkovski and V. Pappa for their technical assistance in mouse genotyping and tissue cutting, R. Grenfell for his technical assistance with the CyTOF acquisition, and T. Adejumo (Fluidigm) for his contribution towards the development of the CyTOF antibody panel, processing of data, and analysis.

This research was supported by Fondazione Italiana Sclerosi Multipla FIMS and Italian Multiple Sclerosis Association AISM Senior research fellowship financed or co financed with the ‘5 per mille’ public funding cod. 2017/B/5 (LPJ); Wellcome Trust Clinical Research Career Development Fellowship G105713 (LPJ); Isaac Newton Trust Research Grant RG 97440 (SP and LPJ); Fondazione Italiana Sclerosi Multipla FIMS and Italian Multiple Sclerosis Association AISM 2018/R/14 (SP and LPJ); National MS Society Research Grant RG 1802-30200 (SP and LPJ); Evelyn Trust (SP); Bascule Charitable Trust (SP); NIHR Cambridge BRC (SP); Medical Research Council UK (MC_UU_00028/4), and Wellcome Trust Investigator award (220257/Z/20/Z) (MPM); Henry Wellcome Fellowship 215888/Z/19/Z (AEV); National MS Society Post-doctoral fellowship FG-2008-36954 (CMW); US National Institute of General and Medical Sciences RM1GM131968 (AD); US National Heart, Lung and Blood Institute R01HL146442, R01HL149714, R01HL148151, R01HL161004, and R21HL150032 (AD); European Committee for Treatment and Research in Multiple Sclerosis (ECTRIMS) Postdoctoral Research Fellowship Exchange Program (G104956) (AMN); Erasmus+ student internship (AVB, LR).

## Author contributions

LPJ, conceptualisation, methodology, *ex vivo* experiments, funding acquisition, supervision, writing; CMW, methodology, *ex vivo* experiments, processing of samples for CyTOF, writing; RH, methodology, ScRNAseq and CyTOF analysis, writing; GK, methodology, *ex vivo* and *in vitro* experiments, processing of samples for CyTOF, tissue pathology, writing; JAH, LC-MS acquisition and data analysis; HAP, MitoP/B acquisition and data analysis; VW, LD-REIMS acquisition and data analysis; YX, LD-REIMS data analysis; AMRVDB, *ex vivo* metabolic flux experiments; AMN, *ex vivo* experiments and *in vitro* FACS; LR, tissue pathology; GB, MitoP/B acquisition; HH, LD-REIMS data analysis; AEV, conceptualisation, methodology; CF, conceptualisation, methodology, writing; CV, conceptualisation, methodology, writing; JCM, conceptualisation, methodology, funding acquisition, supervision; AD, conceptualisation, methodology, funding acquisition, supervision; ZT, conceptualisation, methodology, funding acquisition, supervision; MPM, conceptualisation, methodology, funding acquisition, supervision, writing; SP, conceptualisation, methodology, funding acquisition, supervision, writing.

## Competing interests

SP is founder, CSO and shareholder (>5%) of CITC Ltd and Chair of the Scientific Advisory Board at ReNeuron plc. MPM and HAP have applied for patents in the use of malonate esters to decrease RET in therapeutic situations. Though unrelated to the contents of this manuscript, AD is a founder of Omix Technologies Inc., a founder of Altis Biosciences LLC., scientific advisory board member for Hemanext Inc. and Forma Inc, and a consultant for Rubius Inc. The other authors declare that they have no competing interests.

## Extended Data Figures

**Extended Data Fig. 1.**
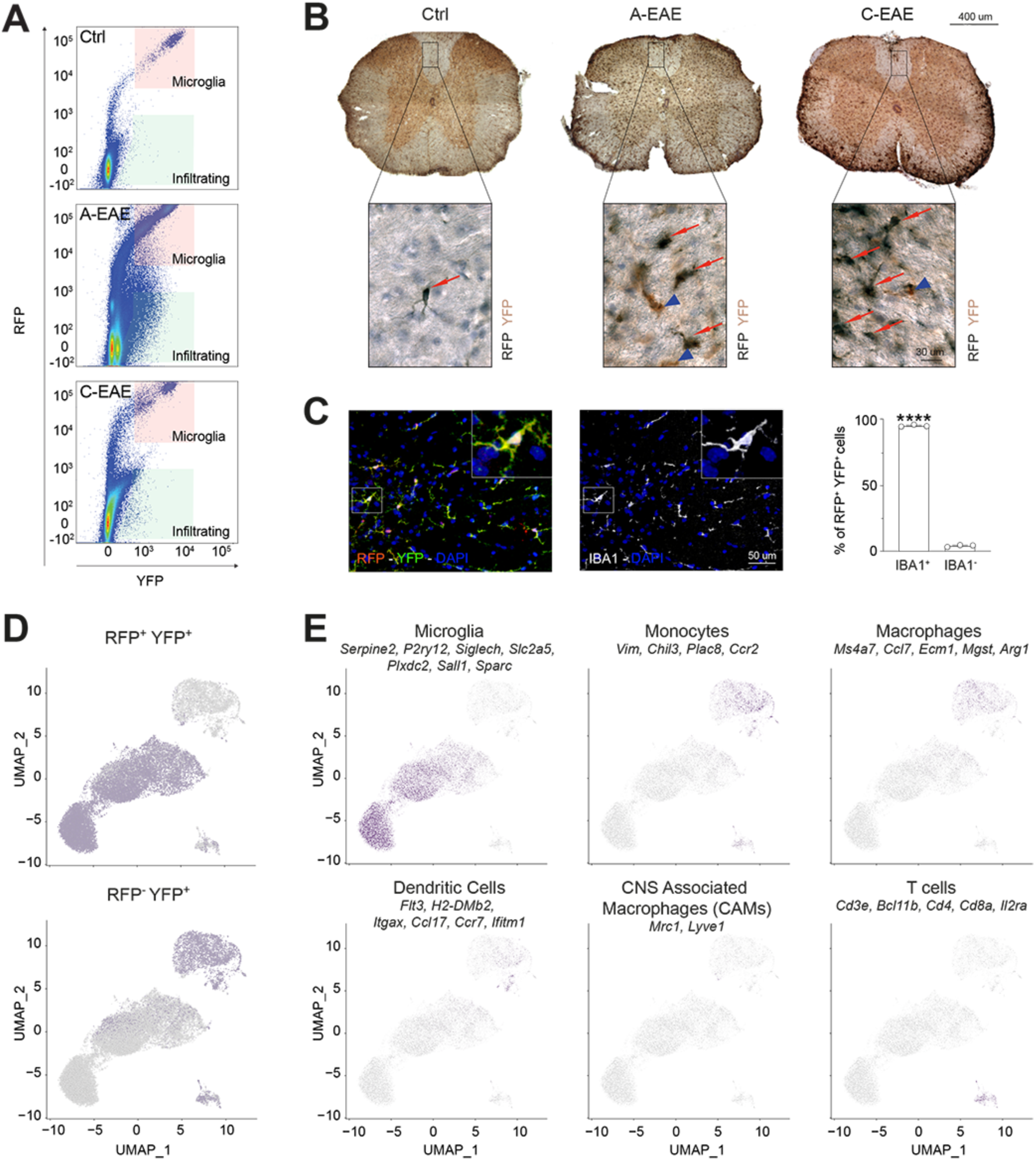
Characterization of microglia and infiltrating myeloid cells in the *Cx3cr1*YFP^CreERT2^:R26^tdTomato^ fate mapping mouse. (**A**) Representative flow cytometry plots of RFP^+^YFP^+^ microglia and RFP^-^YFP^+^ infiltrating myeloid cells in Ctrl, A-EAE, and C-EAE mice. (**B**) Representative immunohistochemistry of RFP^+^YFP^+^ microglia (black, red arrows) and RFP^-^YFP^+^ infiltrating myeloid cells (brown, blue arrowheads) in Ctrl, A-EAE, and C-EAE mice. (**C**) Representative confocal images and quantification of RFP localization showing that 95.87 % (±0.45 SEM) of RFP^+^YFP^+^ microglia co-expressed the myeloid marker IBA1 in *Cx3cr1*YFP^CreERT2^:R26^tdTomato^ mice. ****P<0.0001 (unpaired t-test). (**D**) UMAP plots from the ScRNAseq data with cells (dots) coloured as RFP^+^YFP^+^ microglia and RFP^-^YFP^+^ infiltrating myeloid cells. (**E**) UMAP plots with cells coloured by the sum of the normalized counts of core signature genes for microglia, monocytes, macrophages, dendritic cells, CNS associated macrophages (CAMs), and T cells.

**Extended Data Fig. 2.**
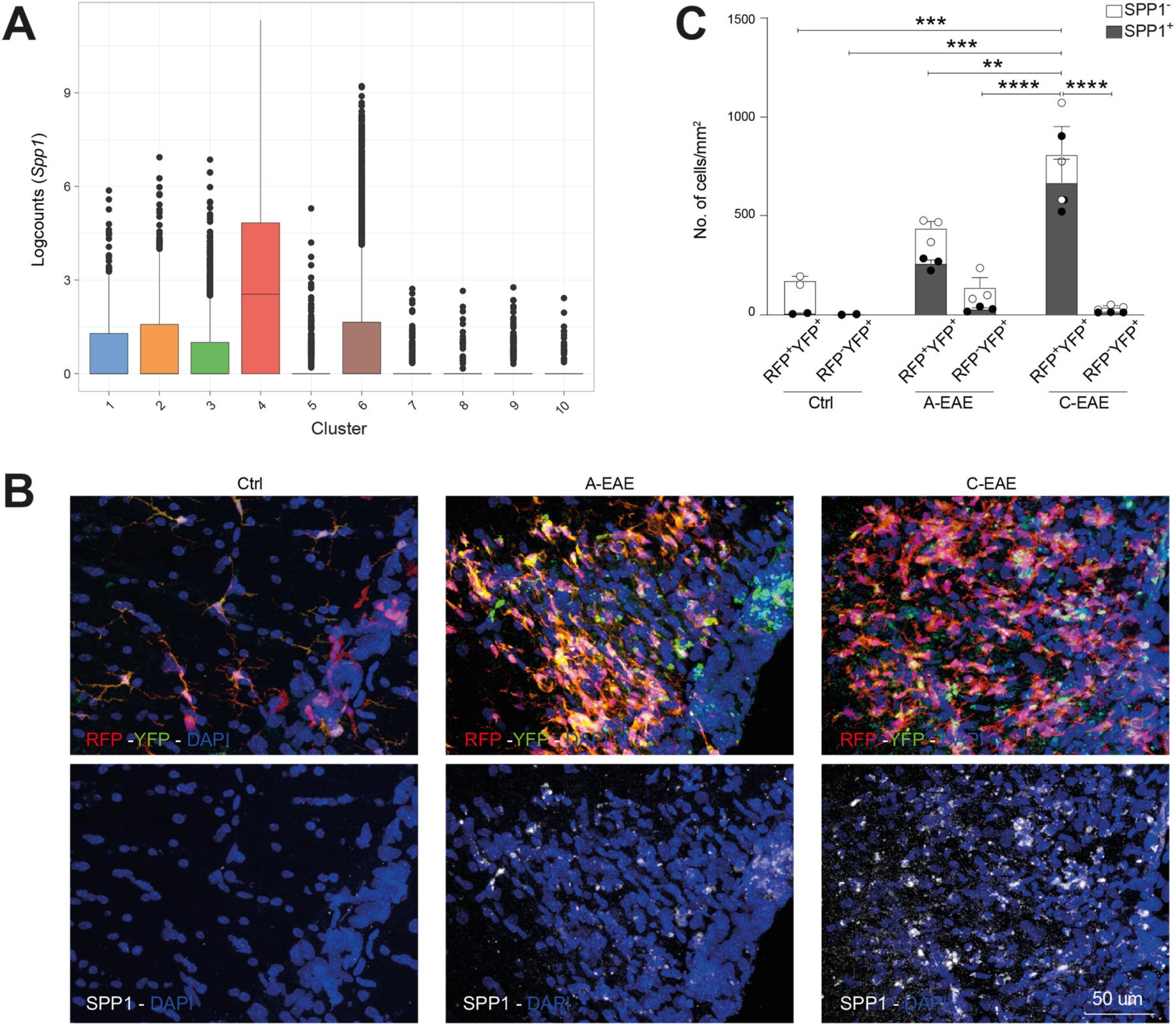
Cluster 4 DAM in *Cx3cr1*YFP^CreERT2^:R26^tdTomato^ Ctrl and EAE mice. (**A**) Box plots of the log_2_-transformed counts of *Spp1* showing its expression in Cluster 4 DAM compared to the other scRNAseq clusters. (**B**) Representative confocal images of SPP1 expression in the posterior columns of the spinal cord in Ctrl, A-EAE, and C-EAE mice. (**C**) Quantification showing the number of SPP1^+^ and SPP1^-^ microglia (RFP^+^YFP^+^) and infiltrating myeloid cells (RFP^-^YFP^+^). Data show that 3.75% of microglia are SPP1^+^ in Ctrl *vs* 58.93% in A-EAE and 82.57% in C-EAE. **P<0.01, ***P<0.001, ****P<0.0001 (one-way ANOVA on SPP1^+^ cells).

**Extended Data Fig. 3.**
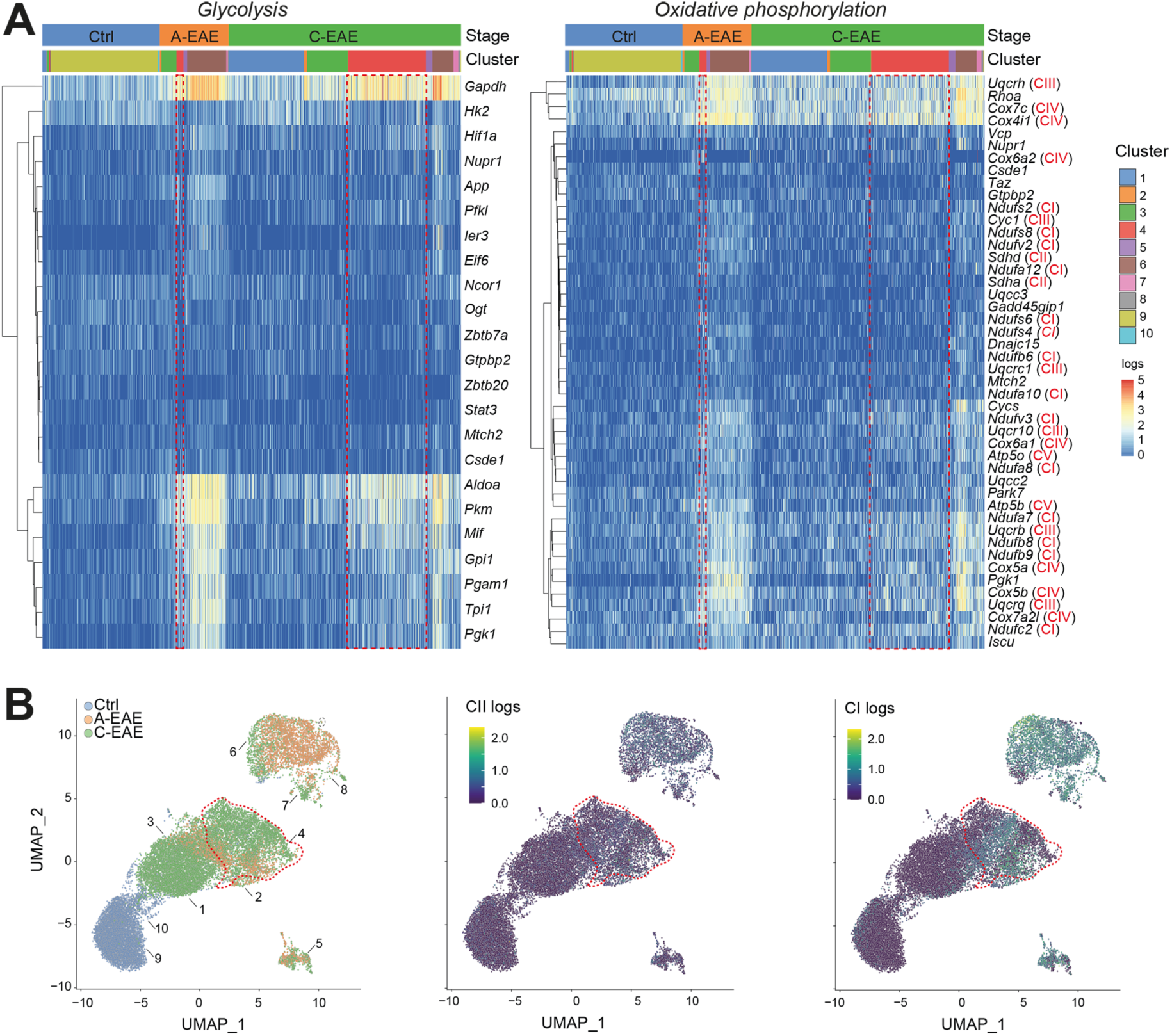
Glycolysis and oxidative phosphorylation complexes genes in myeloid cells isolated from *Cx3cr1*YFP^CreERT2^:R26^tdTomato^ Ctrl and EAE mice. (**A**) Heatmap of the log_2_-transformed counts of the significant differentially expressed genes (rows) involved in glycolysis and oxidative phosphorylation. Red text indicates structural genes of mitochondrial complex I (CI), II (CII), III (CIII), IV (CIV), and V (CV). Dotted red boxes highlight Cluster 4 data. (**B**) ScRNAseq UMAP plots of the whole scRNAseq dataset colored by the stage of disease (left) and the scaled median log_2_-transformed counts of CII (middle) and CI (right).

**Extended Data Fig. 4.**
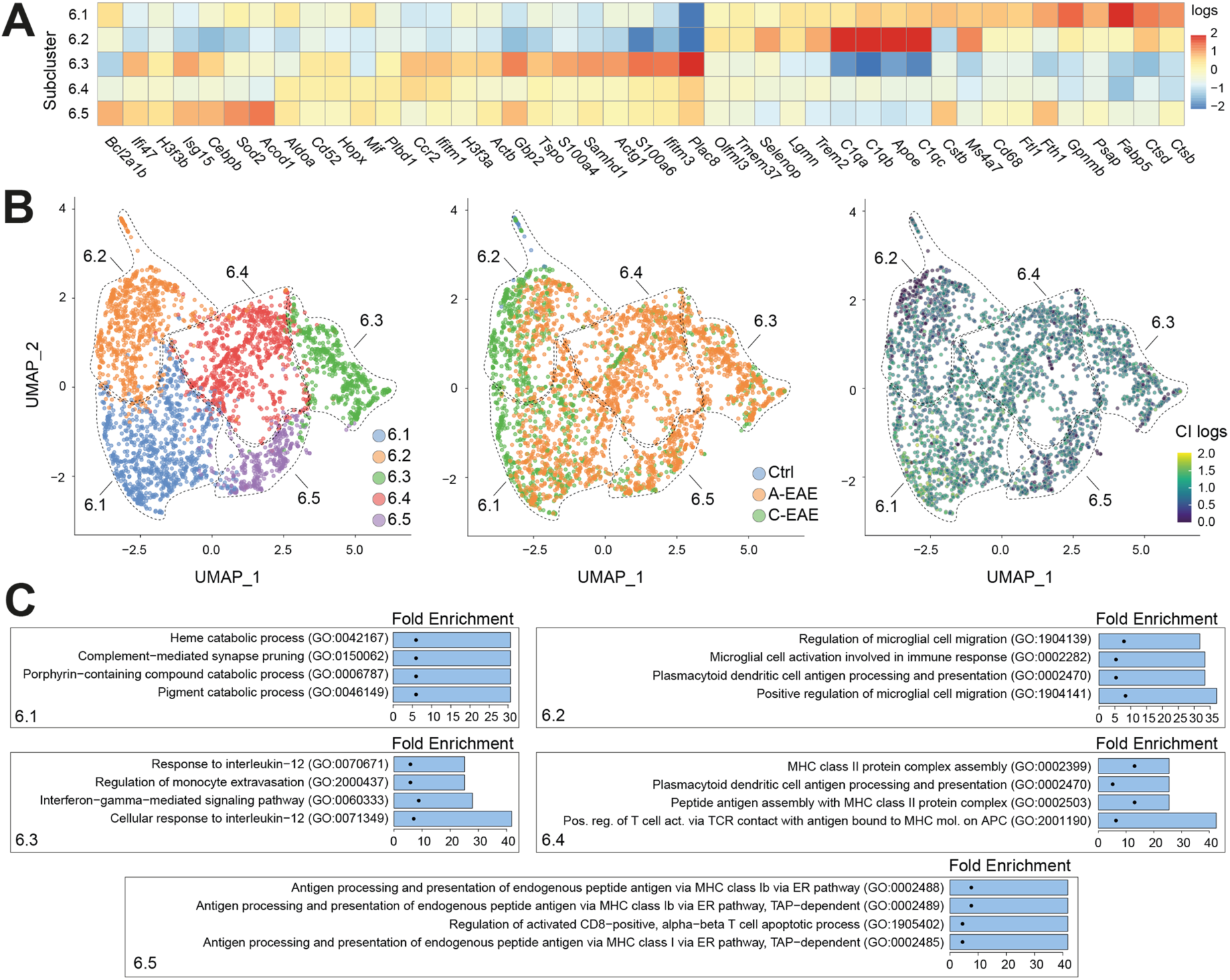
ScRNAseq unsupervised subcluster analysis of Cluster 6 infiltrating myeloid cells isolated from *Cx3cr1*YFP^CreERT2^:R26^tdTomato^ Ctrl and EAE mice. (**A**) Grouped heatmap of the average log_2_-transformed counts of the top differentially expressed genes for the 5 identified subclusters of Cluster 6. (**B**) ScRNAseq UMAP plots of the subclusters of Cluster 6 coloured by subcluster (left), condition (middle), and log_2_-transformed counts of CI (right). (**C**) Top 4 GO terms of the subclusters of Cluster 6.

**Extended Data Fig. 5.**
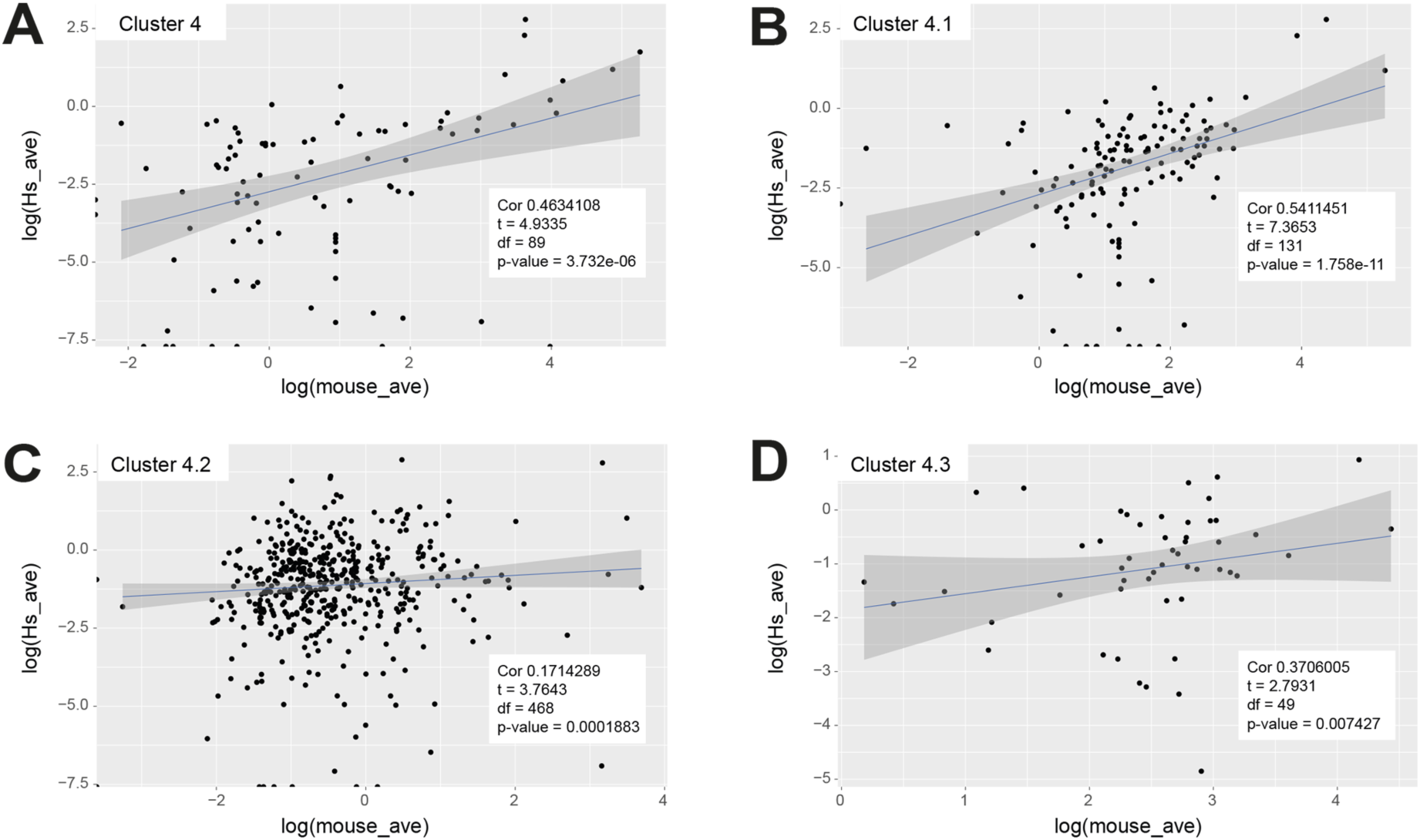
Human microglia activated in progressive MS (MAMS) display a transcriptional profile reminiscent of mouse Cluster 4 DAM. (**A**) Scatter plot depicting the average expression of the upregulated genes from the mouse Cluster 4 DAM, (**B**) Subcluster 4.1, (**C**) Subcluster 4.2, and (**D**) Subcluster 4.3 in the respective mouse cluster *vs* MAMS. The expression across the mouse and human populations was significantly correlated.

**Extended Data Fig. 6.**
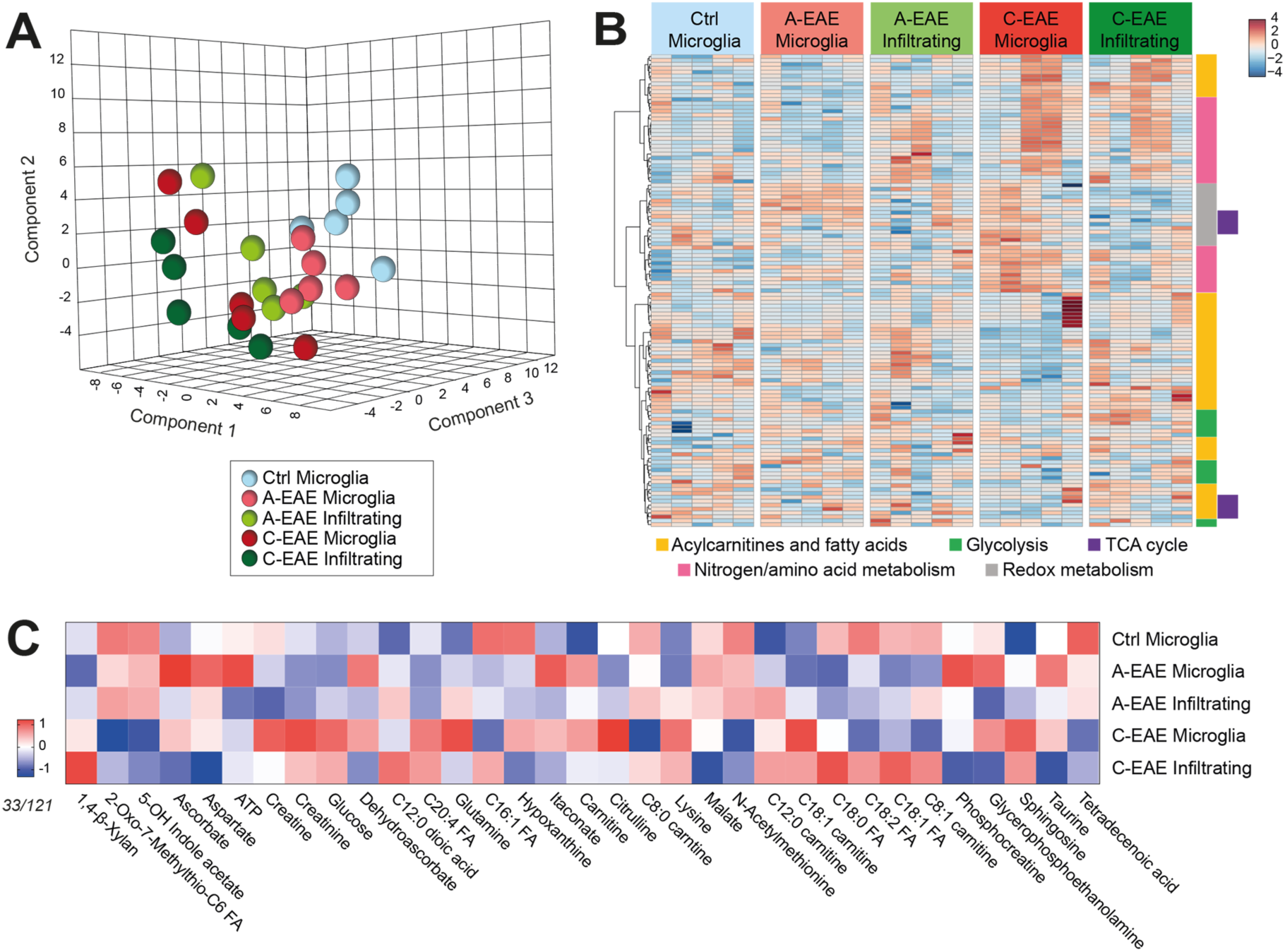
Intracellular LC-MS analysis of microglia and infiltrating myeloid cells isolated from *Cx3cr1*YFP^CreERT2^:R26^tdTomato^ Ctrl and EAE mice. **(A)** Partial least squares discriminant analysis 3D plot of the intracellular metabolome of microglia and infiltrating myeloid cells. **(B)** Heatmap depicting the 125 metabolites (rows) identified via untargeted LC-MS analysis in each biological sample (5 per condition). **(C)** Heatmap showing the 33/121 metabolites found to be significantly differentially regulated when comparing microglia and infiltrating myeloid cells at the different stages of EAE disease (P≤0.05, one-way ANOVA). Scale bars are normalized arbitrary units (A.U.).

**Extended Data Fig. 7.**
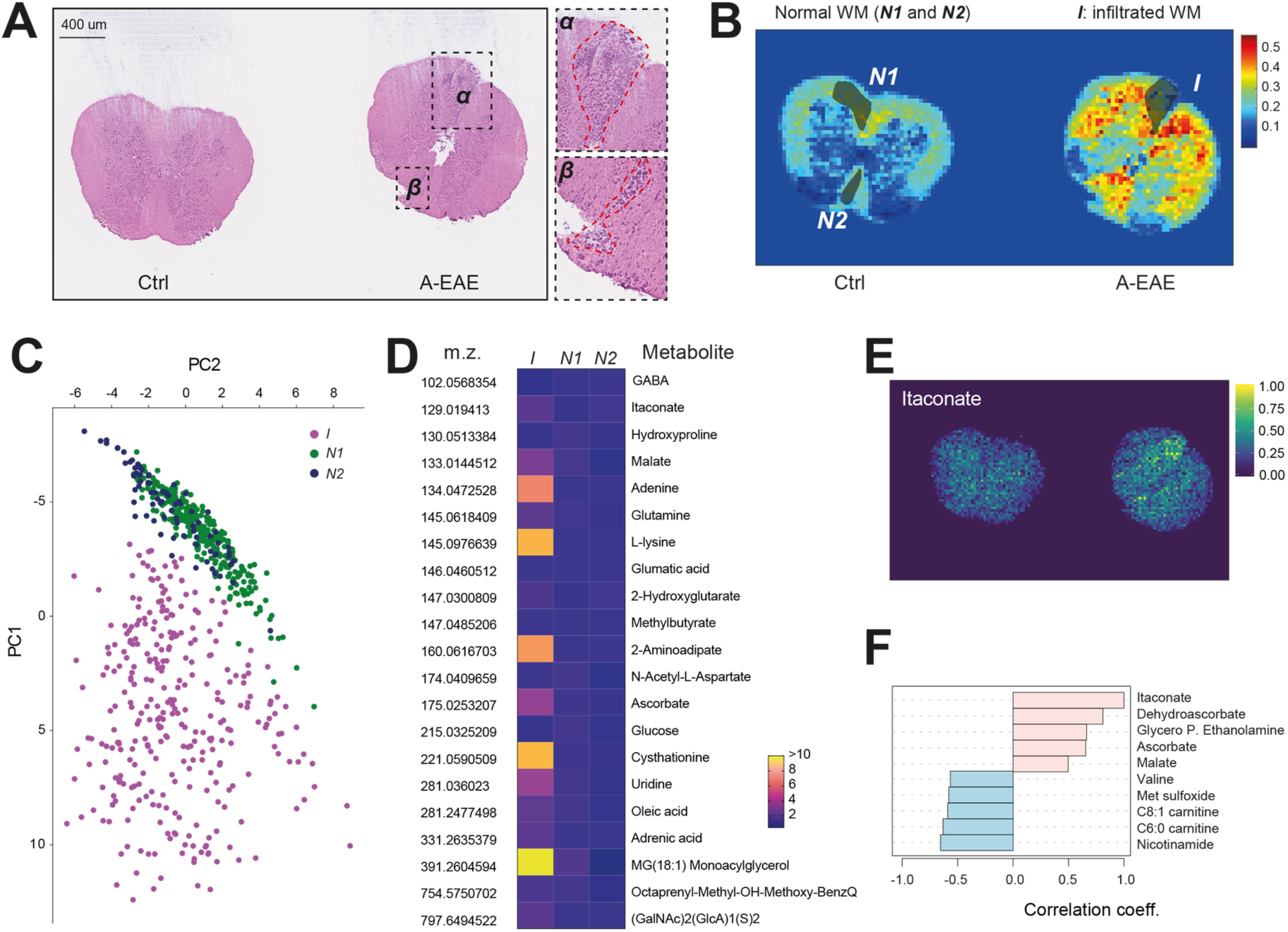
Itaconate levels in Ctrl and A-EAE mice. (**A**) Representative haematoxylin and eosin staining of spinal cord sections from Ctrl and A-EAE mice used for LD-REIMS studies. Magnification of infiltrates in α and β is shown on the right. (**B**) Corresponding colorimetric map based on the multivariate analysis of the spatial metabolome of spinal cord sections, as in A. The infiltrated (*I*) white matter-WM (defined on haematoxylin and eosin staining) was compared with two areas of normal (*N1* and *N2)* WM. Data in the maps are shown as normalized abundances of the (tissue) spectral component extracted from the multivariate analysis. (**C**) PCA plot of the metabolome of *I*, *N1,* and *N2* areas. (**D**) Heatmap showing the metabolites significantly changed between *I*, *N1,* and *N2* areas. Data are shown as fold change over the average values of *N1* and *N*2. **E**) Representative colorimetric maps showing the intensity of the itaconate signal in spinal cord sections as in A. Data in the maps are shown as normalized abundances. **F**) Correlation analysis of itaconate with other metabolites (top 5 up/down) from the whole LC-MS dataset. P.: phosphate.

**Extended Data Fig. 8.**
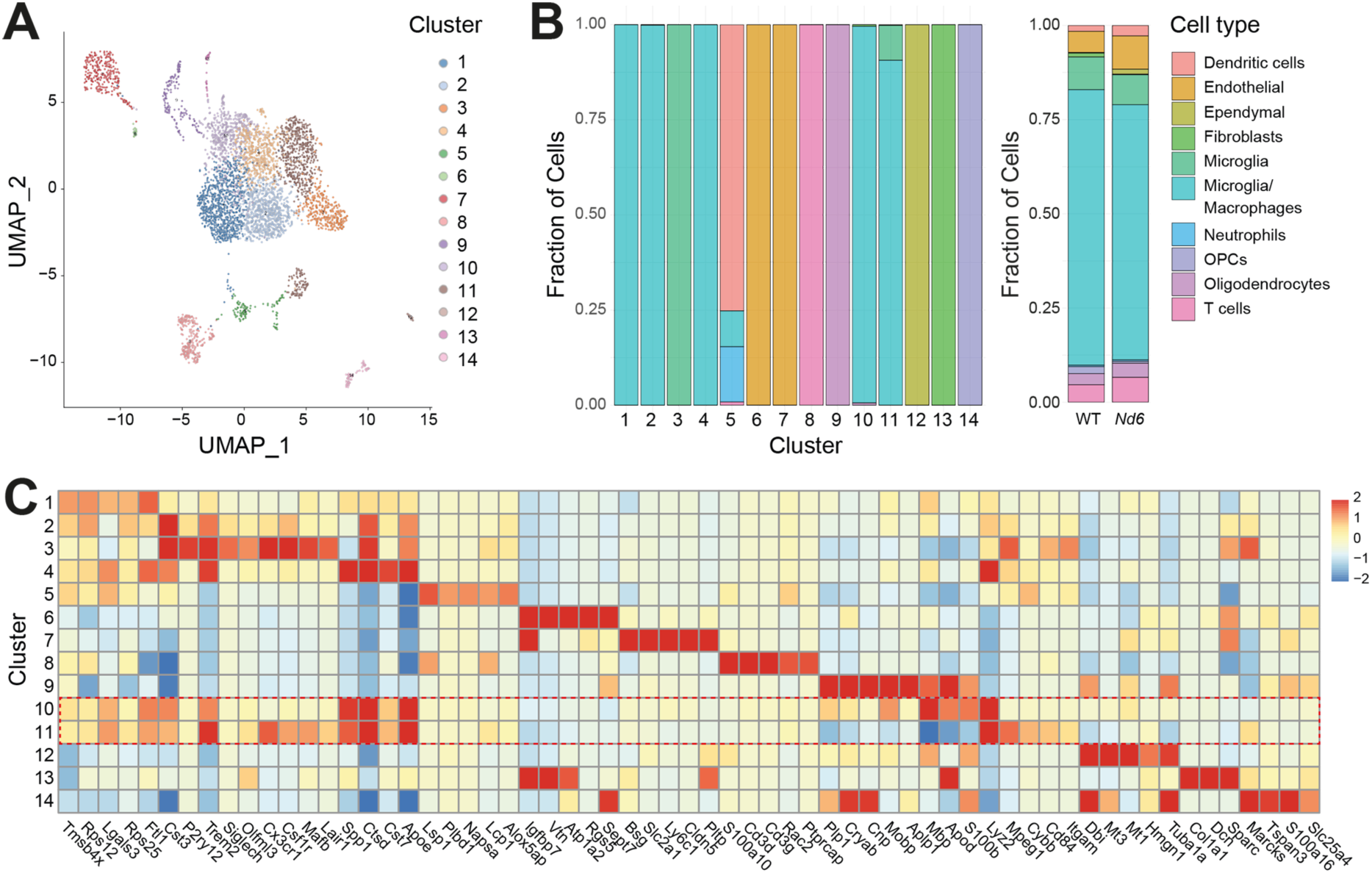
ScRNAseq characterization of the CNS of WT and *Nd6* EAE mice. (**A**) UMAP plot coloured by cluster in WT and *Nd6* EAE mice at 30 dpi. (**B**) Relative proportion of cell types within the 14 clusters (left) and relative proportion of cell types in WT and *Nd6* EAE mice (right). OPCs: oligodendrocyte precursor cells (**C**) Heatmap of the average log2-transformed counts of top 5 differentially expressed genes across the 14 clusters. Dotted red boxes highlight Cluster 10 and 11 DAM.

**Extended Data Fig. 9.**
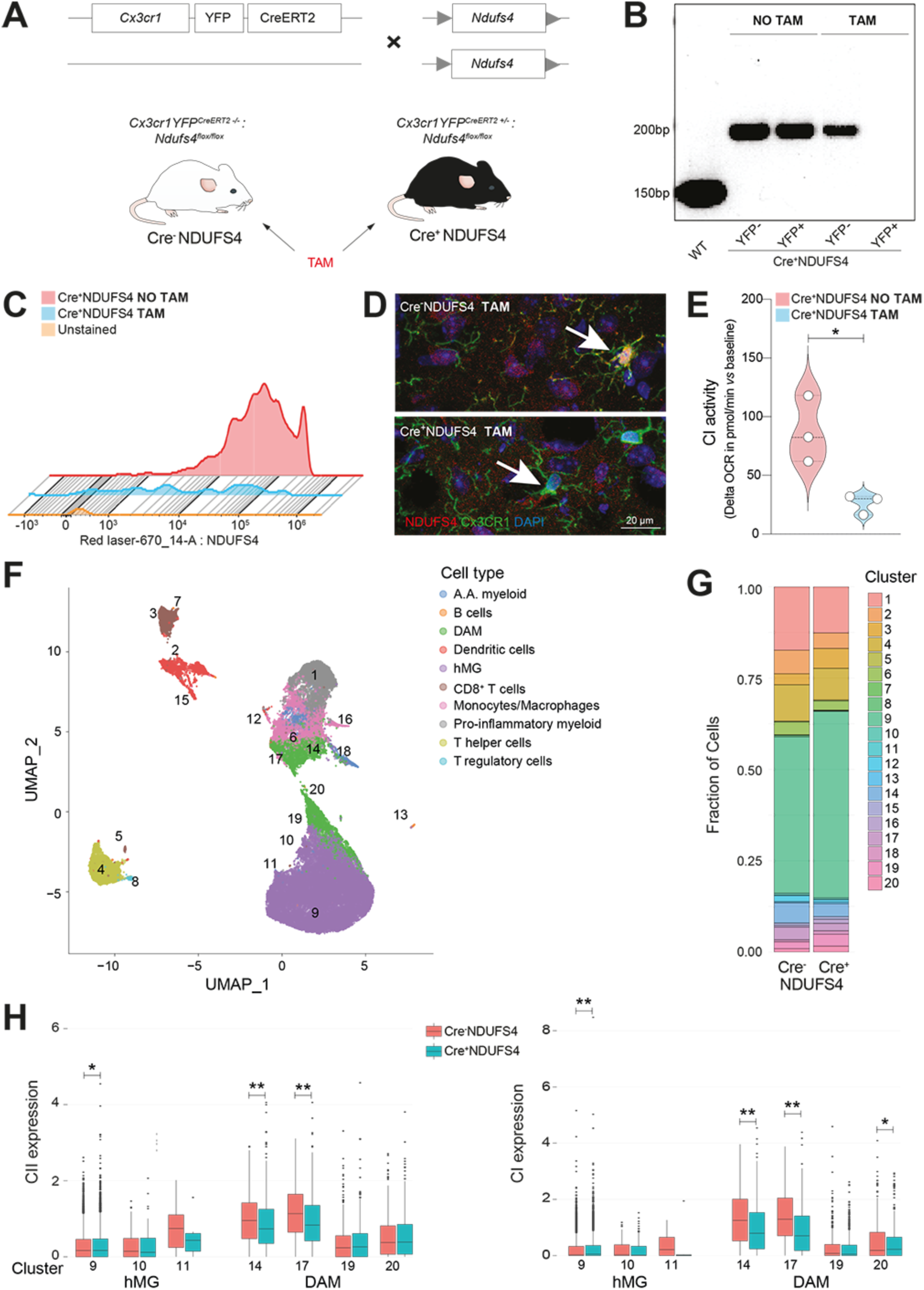
Characterization of the *Cx3cr1*YFP^CreERT2^*:Ndufs4*^flox/flox^ mice and CyTOF from EAE. (**A**) Experimental paradigm for the *in vivo* and *ex vivo* experiments on *Cx3cr1*YFP^CreERT2-/-^*:Ndufs4*^flox/flox^ (Cre^-^NDUFS4) and *Cx3cr1YFP*^CreERT2+/-^*:Nduf4*^flox/flox^ (Cre^+^NDUFS4). A 5-day course of intraperitoneal TAM injections is used to induce transgene activation. (**B**) Genomic PCR for *Ndufs4* from CX3CR1^+^ myeloid cells isolated via FACS from the CNS of healthy Cre^+^NDUFS4 mice 2 days after TAM or saline treatment (NO TAM). (**C**) Flow cytometry showing the NDUFS4 protein expression in CX3CR1^+^ myeloid cells obtained from the CNS of healthy Cre^+^NDUFS4 mice 30 days after TAM or NO TAM. (**D**) Confocal images of NDUFS4 protein expression in CX3CR1^+^ myeloid cells of Cre^-^NDUFS4 and Cre^+^NDUFS4 healthy mice 30 days after TAM. (**E**) CI activity in Cre^+^NDUFS4 EAE mice. TAM treated Cre^+^NDUFS4 EAE mice showed a 70% reduction of CI activity in CX3CR1^+^ myeloid cells at C-EAE compared to Cre^+^NDUFS4 EAE mice NO TAM. *P<0.05 (unpaired t-test). (**F**) UMAP plot from the CyTOF data coloured by cell type found in Cre^-^NDUFS4 and Cre^+^NDUFS4 EAE mice at 50 dpi. Numbers of clusters are shown in superimposition. A.A.: alternatively activated. (**G**) Fraction of cell clusters in Cre^-^NDUFS4 and Cre^+^NDUFS4 EAE mice at 50 dpi. (**H**) CII and CI expression in the clusters identified as hMG and DAM. *P<0.05, **P<0.01 (unpaired t-test).

**Extended Data Fig. 10.**
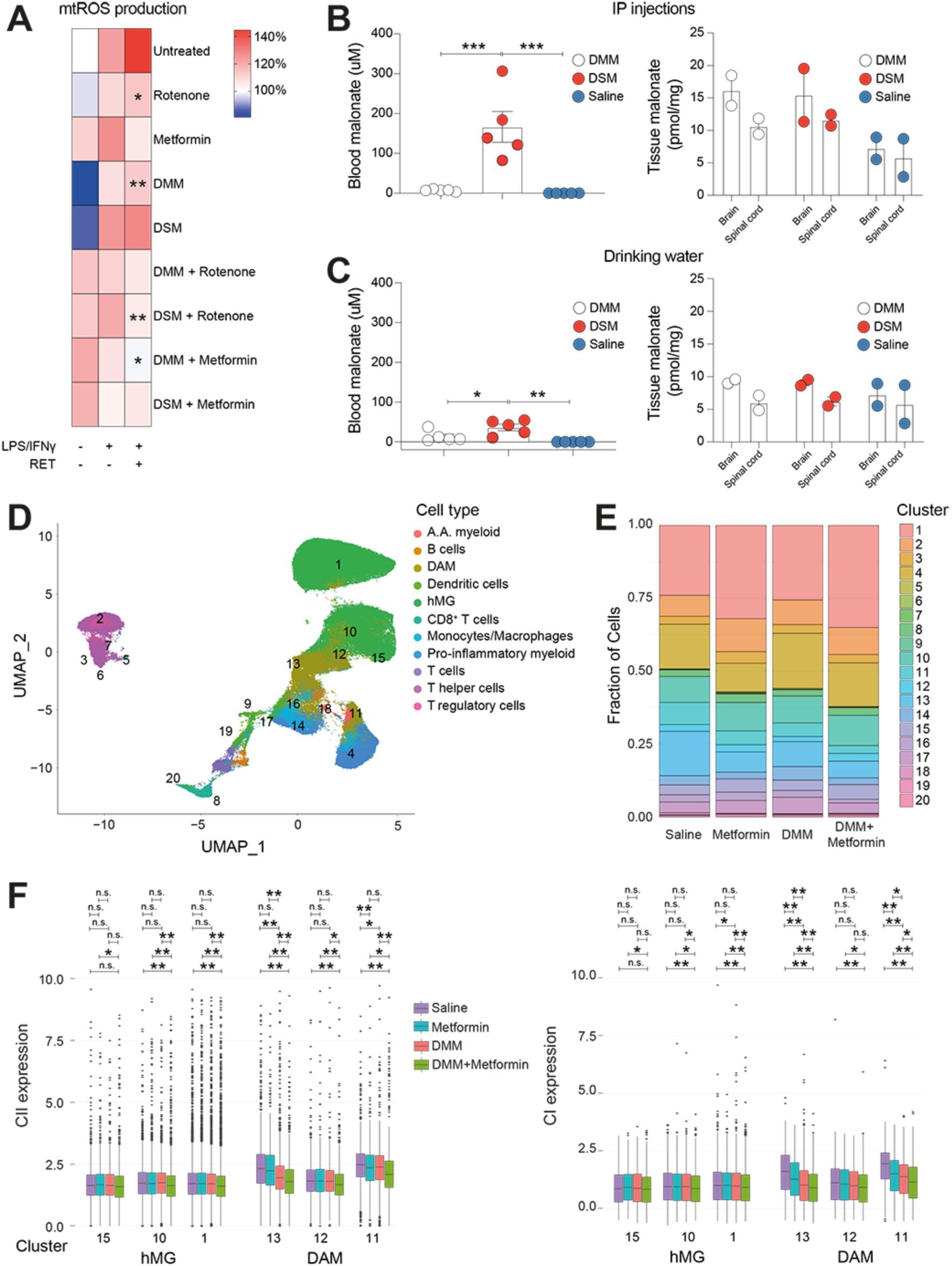
Testing of CII-CI inhibitors and CyTOF from EAE. (**A**) *In vitro* testing of selected CII and CI inhibitors on mtROS production (assessed via MitoSOX) from LPS/IFNγ stimulated microglial cells after RET induction. Data are shown as % increase of mtROS production *vs* unstimulated, untreated cells. *P<0.05, **P<0.01 *vs* untreated RET^+^ pro-inflammatory microglia (one-way ANOVA). (**B-C**) *In vivo* testing of DMM and disodium malonate (DSM) in EAE. (B) EAE mice receiving daily intraperitoneal injections (IP) of DMM (160mg/kg), DSM (160mg/kg), or saline at 7 days from disease onset. Malonate levels in the peripheral blood (at 30 minutes from the daily injections, left) and in the CNS (at sacrifice, right) are shown. (C) EAE mice receiving DMM (1.5%), DSM (1.5%), or saline dissolved in their drinking water (at 7 days from disease onset). Malonate levels in the peripheral blood during treatment (daily, left) and in the CNS (at sacrifice, right) are shown. *P<0.05, **P<0.01, ***P<0.001 (one-way ANOVA). (**D**) UMAP plot from the CyTOF data coloured by cell type found in EAE mice treated with CII and CI inhibitors at 30 dpi. Numbers of clusters are shown in superimposition. A.A.: alternatively activated. (**E**) Fraction of cell clusters in EAE mice treated with Metformin, DMM, DMM+Metformin, and saline-treated controls at 30 dpi. (**F**) CII and CI expression in the CyTOF clusters identified as hMG and DAM. *P<0.01, **P<0.0001 (one-way ANOVA).

## Supplementary Methods

### EAE induction and behaviour

Body weight and EAE clinical score (0= healthy; 1= limp tail; 2= ataxia and/or paresis of hindlimbs; 3= paralysis of hindlimbs and/or paresis of forelimbs; 4= quadriplegia; 5= moribund or death) were recorded daily. At 7-16 days post immunisation (dpi), mice developed the first clinical signs of disease (EAE onset), and at 3 days after disease onset they reached the acute phase of disease (A-EAE). For experiments in the *Cx3cr1*YFP^CreERT2^:R26^tdTomato^ and the *Cx3cr1*YFP^CreERT2^:*Ndufs4*^flox/flox^ mouse lines, EAE mice were kept until 50 dpi to reach the chronic stage of EAE (C-EAE). For *Nd6* and WT mice, EAE was induced and mice were kept until 30 dpi.

### Single-cell RNA sequencing (scRNAseq)

After perfusion, spinal cords were extracted from the spinal columns using a 5 mL syringe filled with ice-cold artificial CSF (aCSF). Spinal cords were mechanically dissociated in a glass Dounce tissue homogenizer with 6 mL of homogenization buffer [aCSF plus 10 mM HEPES (Sigma), 1% bovine serum albumin (BSA) (Sigma), 1 mM EDTA (Thermo Fisher Scientific), 10 mg/ml of DNAse (3000U, Roche), and 40 units/μL of RNAse inhibitor (Invitrogen)]. After tissue dissociation, the suspension was filtered through a pre-wet 40 μM strainer and the homogenizer rinsed with 2 mL of homogenization buffer. Samples were then transferred to 15 mL Falcon tubes and 2.7 ml of 90% Percoll (GE Healthcare) in 10x PBS (Thermo Fisher Scientific) was added to each sample to remove myelin and debris. The 15 mL Falcon tubes were then inverted 10 times gently and the samples were centrifuged at 800 × *g* for 20 minutes at 4°C with a brake speed of 0. Myelin debris visibly layered at the surface was carefully removed with a pipette. Ice-cold buffer [95% autoMACS rinsing solution (Miltenyi Biotec) and 5% MACS BSA (Miltenyi Biotec)] was added to the samples (to fill the 15 mL tubes) and samples were centrifuged at 800 × *g* for 5 minutes at 4°C to remove the remaining Percoll. Pellets were resuspended in ice-cold aCSF (1 mL+7 mL) and then further centrifuged at 800 × *g* for 5 minutes at 4°C. Pelleted cells were resuspended in 200 μL of FACS buffer [Cell Staining Buffer (Biolegend) plus 10 mM HEPES, 1% BSA, 1 mM EDTA, 10 mg/mL of DNAse, and 40 units/μL of RNAse inhibitor] and 7-AAD live/dead stain (Thermo Fisher Scientific) added at a concentration of 1:50. The samples were sorted using a BD FACS Aria III cell sorter set to 3-way purity with a 100 μm nozzle at 20 psi. The live cells gates were set based on the unstained WT sample, 7-AAD stained WT sample treated with DMSO (positive control for cell death), and non-TAM treated *Cx3cr1*YFP^CreERT2^: R26^tdTomato^ samples (to control for bleed-through).

FACS isolated cells were sequenced with single-cell resolution using the v3 10x Genomics Chromium Single Cell 3’ Solution. Up to 18,000 cells per sample were loaded into each well and the resultant libraries were sequenced with the NovaSeq 6000 (Illumina) to depth of at least 50,000 reads per cell as calculated by Cell Ranger. Sequenced samples were demultiplexed and aligned to the GRCm38 (mm10) reference transcriptome using the Cell Ranger 3.1.0 pipeline (10x Genomics). ScRNAseq data processing (post-Cell Ranger) was performed in R version 4.1.3. All packages are publicly available at the Comprehensive R Archive Network (https://cran.r-project.org), the Bioconductor project (http://bioconductor.org), or their respective GitHub pages. The original code is available at https://github.com/regan-hamel/ComplexI.

For each sample, the filtered feature-barcode matrix was loaded and low-quality cells were removed using fixed thresholds of 800 UMIs and 400 unique genes per cell or more than 5% mitochondrial transcripts ^1^. Samples in which the majority of cells did not meet these thresholds were considered failed runs and were not included for downstream analysis.

Likely doublets were predicted and removed using the *cxds_bcds_hybrid()* function from the *scds* package (version 1.2.0) with the parameter *estNdb*l = TRUE ^2^. Briefly, this approach predicts doublets by combining the outputs of two scoring methods: *cxds,* which evaluates the co-expression of genes across cells, looking for instances of unlikely combinations; and *bcds*, which generates artificial doubles based on the dataset, then looks for cell libraries that are similar to the artificial doublets.

Samples were then concatenated, and multi-batch normalization (*batchelor*) was performed to account for differences in sequencing depth across batches. The technical noise (*scran*) was modelled and the top 40% of genes with highest biological variance selected to be used as highly variable genes (HVG) in downstream processing. Correction for technical batch effects was achieved using the mutual nearest neighbours (MNN) approach^3^ by calling *fastMNN()* from the *batchelor* package with *batch* = the Illumina flow cell batch. The batch corrected dimensions were used to visualize the entire dataset in 2D using Uniform Manifold Approximation and Projection (UMAP) (*scater*) with the default parameters.

Cell types were first automatically annotated via the *cellassign* package (version 0.99.21) using a custom marker gene list (Microglia: *Sparc, C1qa, Plxdc2, Serpine2, P2ry12, Tmem119, Siglech, Ctss, Cst3, Slc2a5, Sall1*; Macrophages: *Ms4a7, Ecm1, Arg1, Ccl7, Mgst1*; Monocytes: *Chil3, Plac8, Ccr2, Rgs1, Fn1*; Dendritic Cells: *Cd74, H2-Eb1, H2-Aa, H2-Ab1, Plac8, H2-DMb1, H2-DMa, Klrd1, Flt3, Zbtb46;* Neutrophils: *S100a8, S100a9, Lcn2, Dedd2, Retnlg, Wfdc21, Mmp9;* CAMs: *Ms4a7, Cd74, Cd163, Cbr2, Lyve1, Mrc1;* T-Cells: *Cd2, Cd3d, Cd3e, Cd3g, Cd28, Cd4, Ptprc).* For the *Cx3cr1*YFP^CreERT2^:R26^tdTomato^ dataset, cell labels were then manually adjusted based on expert knowledge and fate mapping labels.

Dataset clustering was achieved by building a shared nearest neighbour graph (*scater*) in the integrated 50 dimensional space with *k = 20* and used the louvain algorithm (*igraph*) to identify clusters. Differentially expressed genes (DEGs) for each cluster were generated by calling the scran function, *findMarkers* with the parameters *group* = cluster membership, *direction* = upregulated, *pval.type* = some, *test.type* = binomial. Clusters of interest were extracted, the HVGs were recalculated for batch correction and the new integrated dimensions for the cluster of interest were used for (sub)clustering, as described above.

DEGs were used for Gene Ontology (GO) enrichment analysis. To this end, the top 600 DEGs for each cluster were put into the PANTHER Classification System overrepresentation test (release 2021-07-02) following the instructions on the Gene Enrichment analysis page (http://geneontology.org/docs/go-enrichment-analysis/), including the optional use of a custom reference list. P-values computed using the default setting of Fisher’s Exact Test were corrected for multiple testing using the Benjamini-Hochberg method. Only biological processes that were over-represented were examined.

For cross-species analysis, the feature-count matrix and meta data was downloaded from GSE180759. Any cell barcode with < 800 UMIs/nucleus, < 400 unique genes/nucleus or > 1% mitochondrial proportions (indicating incomplete nuclear stripping) was removed. Counts were normalised via scan:: logNormCounts() and the UMAP was built based of the top 30% highly variable genes. To quantify the difference between cell subtypes across species, the mouse genes were converted to their human orthologs and used the DEGs from the mouse clusters of interest to build an area under the curve (AUC) score, using the top ranking 5% of genes for each set. Complete analysis is available at https://github.com/regan-hamel/ComplexI.

### Liquid Chromatography-Mass Spectrometry (LC-MS) analysis

Mice were deeply anaesthetized with intraperitoneal injections (IP) of ketamine/xylazine and culled by cervical dislocation. Spinal cords were extracted from the spinal columns using a 5 mL syringe filled with ice-cold sorting buffer [DMEM no phenol red (Thermo Fisher Scientific) + 25mM HEPES (Sigma-Aldrich) + 5% dialysed FBS (Thermo Fisher Scientific) + 1mM EDTA (Honeywell Fluka) + 1x glutaMAX (Thermo Fisher Scientific)]. Spinal cords were mechanically dissociated in a glass Dounce tissue homogenizer with 7 mL of cold homogenization buffer (sorting buffer + 10 mg/mL of DNAse). After tissue dissociation, the suspension was filtered through a pre-wet 40 μM strainer and the homogenizer rinsed with 2 mL of homogenization buffer. Samples were then transferred to 15 mL Falcon tubes and 2.7 ml of 90% Percoll (GE Healthcare) in 10x PBS (Thermo Fisher Scientific) was added to each sample to remove myelin and debris. The 15 mL Falcon tubes were then inverted 10 times gently and the samples were centrifuged at 800 × *g* for 20 minutes at 4°C with a brake speed of 0. Myelin debris visibly layered at the surface was carefully removed with a pipette. Ice-cold buffer [95% autoMACS rinsing solution (Miltenyi Biotec) and 5% MACS BSA (Miltenyi Biotec)] was added to the samples (to fill the 15 mL tubes) and samples were centrifuged at 800 × *g* for 5 minutes at 4°C to remove the remaining Percoll. Pellets were resuspended in sorting buffer (1 mL+7 mL) and then further centrifuged at 800 × *g* for 5 minutes at 4°C. Pelleted cells were resuspended in 100 μL of sorting buffer and 0.5% SYTOX™ blue dead cell stain (Thermo Fisher Scientific). The samples were sorted using a BD FACS Aria III cell sorter set to yield with an 85 μm nozzle.

The live cells gates were set based on the unstained WT sample, SYTOX™ stained WT sample treated with DMSO (positive control for cell death), and non-TAM treated *Cx3cr1*YFP^CreERT2^:R26^tdTomato^ samples (to control for bleed-through). FACS cells were collected in polypropylene tubes, centrifuged at 800 × *g* for 3 minutes at 4°C, and resuspended in ice-cold PBS 1x (at a concentration of 1 million cells/mL).

An additional 1 mL of PBS 1x was added to each sample and cells were further centrifuged at 800 × *g* for 3 minutes at 4°C. Pelleted cells were then resuspended in cold metabolite extraction buffer (at a concentration of 1 million cells/mL), moved to an autosampler vial and stored at -80°C. When all the samples were collected (prior to LC-MS analysis), autosampler vials were immerged in a methanol dry ice bath twice (10 minutes in and 5 minutes out). A total of 10 μL per sample were pulled together to generate a pulled sample control. All samples were then stored at -80°C for LC-MS analysis.

Metabolite extraction was performed at 1 million cells/mL on frozen cell pellets in ice-cold methanol, acetonitrile, and water (5:3:1) and vortexed for 30 min at 4°C. After a centrifugation (18,000 × *g* for 10 min, 4°C), the supernatants were transferred to pre-chilled autosampler vials (Phenomenex) and stored at -80°C for subsequent liquid chromatography coupled to mass spectrometry (LC-MS) analysis. LC-MS analysis was performed on randomized samples using a Vanquish UHPLC coupled online to a Q Exactive Orbitrap mass spectrometer (Thermo Fisher Scientific) in positive and negative ion modes (separate runs). The LC system was fitted with a Kinetex C18 column (150 mm × 2.1 mm, 1.7 μm; Phenomenex); mobile phases and 5-minute gradients were utilized as previously described in detail ^4^. The acquired spectra (.raw) were converted to .mzXML format using RawConverter and signals were subsequently annotated and integrated using Maven (Princeton University) in conjunction with the KEGG database. Quality control samples and instrument stability were assessed as described previously ^5^.

Analysis of LC-MS data was performed with MetaboAnalyst 5.0 (https://www.metaboanalyst.ca/). Features with >50% missing values were removed (ADP-ribose, IMP, UDP-glucose, UDP-GlcNAc were also removed as there were too many missing values). The remaining missing values were estimated using KNN with no data filtering. Data were normalised by median, followed by Log transformation, and auto scaling to obtain normalised arbitrary units (A.U.) Partial least squares - discriminant analysis (PLS-DA) was performed on the whole LC-MS dataset to generate 3-D scores plot among selected components. A hierarchical clustering heatmap was generated on normalised data using autoscale features and Ward clustering method. Statistical analysis of metabolites found to be significantly altered among all conditions was performed via one-way ANOVA (Graph Pad Prism 9 for macOS, GraphPad Software, San Diego, CA, USA, www.graphpad.com) followed by unpaired t-test for multiple comparisons (Supplementary Data 3). Direct comparisons between cells belonging to two different disease stages were performed to generate volcano plots (fold change threshold: 2.0; P-value threshold: 0.1). Correlation analysis against a given feature or stage was performed with the PatternHunter tool of MetaboAnalyst using Pearson r as distance measure.

### Mitochondrial membrane potential of microglial BV2 cells *in vitro*

BV2 microglial cells (gift from Aviva Tolkovsky) were kept in culture and expanded in expansion medium [DMEM high glucose (GIBCO), 2% FBS (Thermo Fisher Scientific), 1% pen/strep (Thermo Fisher Scientific)]. After reaching 70% confluency, BV2 cells were washed with warm PBS 1x, detached with Trypsin-EDTA 0.05%, (Thermo Fisher Scientific) for 3 minutes followed by trypsin neutralization in expansion media.

BV2 cells were pelleted at 300 × *g* for 5 minutes and seeded with experimental media [no phenol read DMEM high glucose (GIBCO), 1% pen/strep] in a 96-well black clear bottom plate (Thermo Fisher Scientific) at a density of 1×10^5^ cells per well. After 16 hours, BV2 cells were stimulated with LPS 100 ng/mL (serotype EH100, Enzo) plus IFN*γ* 10 ng/mL (Peprotech).

Cells were left for 23 hours, and reverse electron transport (RET) was induced by adding oligomycin at a final concentration of 2 *μ*M and dimethyl succinate at a final concentration of 20 mM (Sigma-Aldrich). After 15 minutes at 37°C, cells were stained with TMRM (final concentration 50 nM) and Rotenone 1 *μ*M (Sigma-Aldrich) was added to wells where complex I was inhibited. After 30 minutes, TMRM signal was read using a yellow laser 582/15 at Ex548/Em575 with a Tecan Infinite M200 Pro plate reader.

### Mitochondrial superoxide production of microglial BV2 cells *in vitro*

BV2 cells were maintained, cultured, and replated in a 96-well black clear bottom plate and RET was induced, as described above. After 15 minutes at 37°C (5% CO_2_ incubator), cells were stained with MitoSOX™ (Thermo Fisher Scientific) (5*η*M) and rotenone 1 *μ*M (Sigma-Aldrich), Metformin 10 mM (Sigma-Aldrich), dimethyl malonate-DMM 10mM (Sigma-Aldrich), or disodium malonate-DSM 10mM (Sigma-Aldrich) were added. After 30 minutes, the MitoSOX™ signal was read using a blue laser 595/40 at Ex510/Em580 with a Tecan Infinite M200 Pro plate reader.

For the analysis of MitoSOX™ via FACS, BV2 cells were seeded at 0.5 × 10^6^ cells/well in a 6-well plate in 1 mL of experimental media [DMEM (Thermo Fisher Scientific) with 1% pen/strep (Thermo Fisher Scientific)]. At 24 hours after seeding, the media was replaced with 1 mL of fresh experimental media. A working solution of LPS (10 *μ*l of stock LPS into 990 *μ*l of 1x PBS) and IFN*γ* (2 *μ*l of 10,000x stock IFN*γ* into 98 *μ*l of 1x PBS) was prepared. BV2 cells were treated for 23 hours with 10 μL of the LPS working solution and 5 *μ*l of the IFN*γ* working solution. 60 minutes before the end of the 24 hrs LPS/IFN*γ* treatment timepoint, RET was induced (as described above). After 30 minutes, the working solution of rotenone was added to the well. At 24 hours from treatment, the media was collected from the well into a 15 mL conical tube. Then, 500 *μ*l of Accutase cell dissociation reagent (ThermoFisher Scientific) was added to the well and the plate incubated at 37°C for 5 minutes. After 5 minutes, 1 mL of 1x PBS was added to the well and the cells were transferred to the 15 mL conical tube with the conditioned media. The cells were then pelleted at 300 × *g* for 5 minutes. The supernatant was removed, and the cell pellet was resuspended in 100 *μ*l of MitoSOX™ Red working solution (1 *μ*l of a 5mM stock into 1 mL of 1x PBS). The cell suspension was moved to a 1.5 mL microcentrifuge tube and incubated at 37°C in an Eppendorf Thermomixer comfort machine set to 500 RPM for 30 minutes protected from light. After 30 minutes, labelled cells were pelleted at 300 × *g* for 5 minutes. The supernatant was removed, and the cell pellet was resuspended in 300 µL of 1x PBS and transferred to a polystyrene flow cytometry tube (Corning). The samples were kept on ice until acquisition. 10 minutes prior to analysis, a working solution of DAPI (1:1000 in 1x PBS) (Millipore Sigma) was added to the sample (1:100 of working solution). Flow cytometry gates were set on DAPI- and MitoSOX™ Red^+^ positive controls and >200,000 total events were captured using a BD FACS Aria III cell analyser. The FCS files were exported, and analysis was performed using FlowJo v10 software (BD Biosciences).

### Microglial BV2 cells and SH-SY5Y *in vitro* co-cultures

SH-SY5Y cells (gift from Michael Whitehead) were kept in culture and expanded in growth medium [DMEM-F12 (Gibco), 10% FBS, 1% pen/strep]. Growth medium was refreshed every 4 to 7 days. After reaching 80-90% confluency, SH-SY5Y cells were washed with warm PBS 1x (Gibco), detached with Trypsin-EDTA 0.05%, (Thermo Fisher Scientific) for 3 minutes followed by trypsin neutralization in growth media and spun down at 0.4 × *g* for 5 min. Next, SH-SY5Y cells were counted and seeded at a density of 5×10^4^ cells/well on coverslips in 12-well plates with differentiation media [Neurobasal medium (Thermo Fisher Scientific) B27 supplement 2% (Thermo Fisher Scientific), GlutaMAX 1% (Thermo Fisher Scientific), and all-trans retinoic acid 10*η*M (STEMCELL Technologies). Differentiation media was replaced every other day until day 9 post-seeding for co-culture experiments with BV2 microglial cells. BV2 cells were maintained and cultured as described above. To time the BV2 and SH-SY5Y co-cultures, on day 7 post-seeding of the SH-SY5Y cells, BV2 cells (70% confluency) were spun at 300 × *g* for 5 min and seeded inside a 0.4 *μ*m-pore size trans well insert (24 well-size, Millipore) at a density of 1×10^5^ cells/trans well with 1 mL of experimental media [DMEM high glucose (GIBCO), 1% pen/strep]. Trans well inserts were placed in a 12-well plate prefilled with 1 mL of experimental media per well. The next day (day 8 post-seeding of the SH-SY5Y cells), BV2 cells were stimulated with LPS 100 ng/mL (serotype EH100, Enzo) plus IFN*γ* 10 ng/mL (Peprotech). After 12 hours from the LPS/IFN*γ* stimulation (day 9 post-seeding of the SH-SY5Y cells), RET was induced by adding oligomycin at a final concentration of 2 *μ*M (Sigma-Aldrich) and dimethyl succinate at a final concentration of 20 mM (Sigma-Aldrich) to the BV2 trans well for 15 minutes at 37°C. Rotenone 1 *μ*M (Sigma-Aldrich) was added to trans well to inhibit complex I. BV2 cells were further incubated for 30 minutes at 37°C. Supernatants from all the trans wells and underlying wells were then removed. Trans wells with BV2 cells were washed with pre-warmed 1x PBS (Gibco) and transferred to a 12-well plate prefilled with 2 mL of 1x PBS. Trans wells with BV2 cells were washed with pre-warmed 1x PBS again, replaced with 1 mL of experimental media, and then put into co-culture with SH-SY5Y cells. After 6 hours of co-culture, trans wells and the media were removed. SH-SY5Y cells were washed with pre-warmed 1x PBS and processed for downstream analyses.

For neurite length analysis, 10x bright field pictures of 5 ROIs per coverslip from each condition were taken using a Leica DM IL LED microscope. Images were converted to 8-bit and contrast was adjusted so the neurites were easily visible using Fiji 2.0.0. software. Data were quantified by semi-automatically tracking NeuronJ plugin from an average of >30 neurite lengths per ROI from n>3 independent experiments.

Catalase gene expression analysis was performed via qRT-PCR. At the end of co-culture experiment, total RNA from SH-SY5Y cells was collected by washing cells with ice-cold 1x PBS and adding 350 μL of RLT lysis buffer (QIAGEN). Samples were then stored at −80°C until extraction. Total RNA from SH-SY5Y was extracted using the RNeasy Mini Kit (QIAGEN) following the manufacturer’s instructions. For qRT-PCR analysis, equal amounts of RNA were reversed-transcribed using the high-capacity cDNA reverse transcription kit (Thermo Fisher Scientific) according to the manufacturer’s instructions. cDNA was then quantified with the NanoDrop 2000c instrument (Thermo Fisher Scientific) and qRT-PCR was performed with the TaqMan™ Fast Universal PCR Master Mix (2x) (Thermo Fisher Scientific) and TaqMan® Gene Expression Assays for: *Catalase* (Thermo Fisher Scientific, Hs00156308_m1) and 18S (Thermo Fisher Scientific) was used for normalization. Samples were run in triplicates using a 7500 Fast Real-Time PCR System (Applied Biosystems) and analysed with the 2^-ΔΔCT^ method.

### Assessment of *ex vivo* ROS production

ROS production was assessed in myeloid cells isolated *ex vivo* from *Cx3cr1*YFP^CreERT2^:R26^tdTomato^ mice via FACS, as described for cells undergoing LC-MS analysis. After FACS, cells were collected in polypropylene tubes, centrifuged at 800 × *g* for 3 minutes at 4°C, and resuspended in 1x PBS (at a concentration of 1 million cells/mL). FACS sorted cells were centrifuged at 800 × g for 3 minutes at 4°C and the pellet was resuspended at 1 million cells/ml in sorting buffer. A 96-well plate was coated with Cell-Tak (Corning) and washed with dH_2_O. Cells were seeded at 10,000 cells per well and the plate was centrifuged at 200 × *g* for 1 minute with 0 brake. The cells were stained using the CellROX™ Deep Red Flow Cytometry Assay Kit (Thermo Fisher Scientific) following the manufacturer’s instructions. Briefly, 100 µL of sorting buffer was added to each stained well, and 50 µl sorting buffer + 50 µl tert-butyl hydroperoxide solution (TBHP, an inducer of ROS) was added to each positive control well. A total of 2 µl of 250 µM CellROX™ Deep Red reagent was added to all wells, and the plate was incubated for 30 minutes at 37°C. The cells were fixed in 4% paraformaldehyde (PFA) (Thermo Fisher Scientific) for 10 minutes and stained with 4’,6-diamidino-2-phenylindole (DAPI). Images were acquired on the Leica microsystems CTR4000 within 2 hours. Using Fiji 2.0.0. software, the intensity of CellROX™ Deep Red over DAPI was calculated. The fold change of ROS over the course of disease was calculated compared to control (non-immunised) mice.

To study mitochondrial ROS production in EAE mice, we used the ratiometric mass spectrometric mitochondria-targeted ROS probe MitoB, as previously described ^6^. This probe is rapidly taken up by mitochondria *in vivo* and then oxidized to MitoP by the ROS hydrogen peroxide and peroxynitrite. Consequently, measuring the MitoP/MitoB ratio by LC-MS/MS indicates changes in mtROS *in vivo*. Mice were placed in a heat box to expose the tail vein. Mice were injected with 100 µl of 0.25 mM MitoB (Cayman Chemical) in saline. Three hours after injection, mice were sacrificed and 30-50 g of the cerebellum, the rest of the brain, and the spinal cord were collected separately and flash frozen in liquid nitrogen. 30-50 mg of tissue was homogenised in a Precellys 24 (Bertin Instruments) using pre-filled bead mill tubes (Thermo Fisher Scientific) in 60% acetonitrile with 0.1% formic acid (6,500 rpm; 15 s × 2 cooling on ice in between). The tissue was then centrifuged at 16,000 × *g* for 10 minutes and the supernatant was collected. 200 µl of 60% acetonitrile with 0.1% formic acid was added to the pellet, vortexed and centrifuged at 16,000 × *g* for 10 minutes. The supernatant was combined. An internal standard of 10 µM d_15_-MitoB and 5 µM d_15_-MitoP was added to the supernatant which was then vortexed and incubated at -20°C for 30 minutes. The sample was centrifuged at 16,000 × *g* for 10 minutes and the supernatant was transferred to a 96 well filter plate. The supernatant was filtered and dried in a speed vac overnight. The pellet was resuspended in 20% acetonitrile with 0.1% formic acid, vortexed for 5 minutes, and centrifuged at 16,000 × *g* for 10 minutes. The samples were then run on the LC-MS/MS for analysis. LC-MS/MS analysis was performed using Waters Xevo TQ-S mass spectrometer (Waters, UK), samples were stored in a cooled autosampler at 10°C and injected at 2 µL. Separation was achieved using an Acquity UPLC BEH 1.7µM C18 Column at 40°C (Waters, UK). MS buffers of A) 95% H_2_O/5% acetonitrile/0.1% formic acid, and B) 90% acetonitrile/10% H_2_O/0.1% formic acid were infused at 200 μL min using this following gradient: 0–0.3 min, 5% B; 0.3–3.0 min, 5–100% B; 3–4 min, 100% B; 4.0–4.1 min, 100–5% B; 4.1–4.6 min, 5% B. Eluant was diverted to waste at 0–1 min and 4–4.6 min. For mass spectrometry, electrospray ionization was used in positive mode with nitrogen as the desolvation gas and argon as the collision gas. Instrument parameters were set to the following, cone voltage, 79 V; source spray voltage, 3.2 kV; ion source temperature 150°C; collision energy, 65V. For quantification, the following transitions were used: MitoB, 397>183; d_15_–MitoB, 412>191; MitoP, 369>183; d_15_–MitoP, 384>191. Spectral data was processed using TargetLynx and all data was calculated based on MS response relevant to the internal standards.

### Metabolic flux analysis

A Seahorse calibration plate was rehydrated with calibration buffer (Agilent) overnight in a non-CO_2_ incubator at 37°C the day before FACS isolation of cells. On the day of FACS, the seahorse cell culture plate was coated with Cell-Tak (Corning) and washed with dH_2_O. After FACS, cells were collected in polypropylene tubes, centrifuged at 800 × *g* for 3 minutes at 4°C, and resuspended in 1x PBS (at a concentration of 1 million cells/mL). Cells were then centrifuged at 800 × *g* for 3 minutes at 4°C and resuspended at 1 million cells/mL in sorting buffer. Cells were seeded on the Seahorse XFe96 plate at 40,000 cells per well (up to six replicates per mouse per cell type) and centrifuged at 200 × *g* for 1 minute with 0 brake. The culture plate was placed in an incubator at 37°C for 30 minutes and then transferred into a non-CO_2_ incubator for 30 minutes. The plate was calibrated, and the cells were washed once with mitochondrial assay solution (MAS, made of sucrose 70mM + mannitol 220 mM + KH_2_PO_4_ 10mM + HEPES 2mM + EGTA 1mM, in dH_2_O at pH=7.2). Just before loading the plate in the Seahorse XFe96 analyser, cells were put in MAS + 0.2% BSA + 1 nM plasma membrane permeabilizer (PMP) (Agilent) + 4 mM ADP (pH 7.4). The cell plate was equilibrated, and the baseline was measured. Glutamate 17.5 mM + malate 17.5 mM were used to activate complex I. Rotenone (3µM, final concentration well) was used to inhibit complex I and disodium succinate (10 mM, final concentration well) (Sigma-Aldrich) was used to activate complex II. Antimycin A (4 µM, final concentration well) was used to inhibit complex III and stop mitochondrial respiration. Five reads were obtained (every 4.63 seconds) at baseline and after each injection. If cells did not respond to the addition of Antimycin A, they were excluded from further analysis. After the Seahorse run, the cells were fixed with 4% PFA for 10 minutes and stained with DAPI for 2 minutes. Using a Leica microsystems CTR4000 to take images at 10x, the OCR values of every plate were normalised based on the mean area of DAPI of four regions of interest. Seahorse analysis was performed using Wave 2.6.1. Complex I activity was calculated as the percentage change decrease of the mean of OCR obtained from 2 consecutive positive measurements before and after rotenone injection. Complex II activity was calculated as the percentage change decrease of the mean of OCR obtained from 2 consecutive positive measurements before and after succinate injection.

### Genomic PCR

Samples were lysed in a thermal cycler: 65°C for 15 minutes, 96°C for 2 minutes, 65°C for 4 minutes, 96°C for 1 minute, 65°C for 1 minute, 96°C for 30 seconds, 4 °C on hold and molecular biology graded water (120 µL , Thermo Fisher Scientific) was added to each sample. A total of 20-100 ng of DNA from each sample was added to a master mix containing molecular biology graded water (7.5 µl) and BioMix Red (12.5 µl, Bioline) with the appropriate amount of primers. Primer sequence for *Cx3cr1*^Cre^ mice *–* C: 5’-AAG ACT CAC GTG GAC CTG CT-3’; mt – R: 5’-CGG TTA TTC AAC TTG CAC CA-3’; wt – R: 5’-AGG ATG TTG ACT TCC GAG TTG-3’; expected product: mutant 300 base pairs (bp), heterozygote 300 bp and 695 pb, wild-type 695 bp. Primer sequence for R26^tdTomato^ mice – mt F: 5’ - CTG TTC CTG TAC GGC ATG G – 3’; mt R: 5’ - GGC ATT AAA GCA GCG TAT CC – 3’; wt F: 5’ - AAG GGA GCT GCA GTG GAG TA – 3’; wt R: 5’ - GGC ATT AAA GCA GCG TAT CC – 3’; expected products: mutant 196 bp, heterozygote 297 bp and 196 bp, wilt-type 297 bp. Primer sequence for *Nd6* mice – F: 5’ - TAC CCG CAA ACA AAG ATC ACC CAG - 3’; R: 5’ - TTA GGC AGA CTC CTA GAA GG - 3’. Primer sequence for *ndufs4* mice – C loxP_A: 5’- AGC CTG TTC TCA TAC CTC GG – 3’; mt R loxP_B: 5’ – GCT CTC TAT GAG GGT ACA GAG – 3’: wt R loxP_C: 5’ – GGT GCA TAC TTA TAC TAC TAG TAG – 3’; expected products: mutant 200 bp, heterozygote 200 bp and 150 bp, wild-type 150 bp. Amplification reaction parameters: initial denaturation at 94°C for 4 minutes (1x); denaturation 94°C for 30 seconds, annealing temperature 65°C for 30 seconds, elongation time (ET) 72°C for 40 seconds (repeated 35x); final extension 72°C for 8 minutes, and 4°C on hold. The same PCR protocol was applied for the genomic analysis of cells isolated via FACS from *Cx3cr1*YFP^CreERT2^:*Ndufs4*^flox/flox^ mice, where DNA was extracted using the DNeasy Blood & Tissue Kit (Qiagen) following the manufacturer’s instruction. Next, amplified nucleic acids (1 μL) were mixed with master mix (24 μL, see above), and loaded onto a 3% agarose gel [agarose (Bioline), tris buffered saline TAE (Sigma-Aldrich), red nucleic acid gel stain (Biotium)] with a TrackIt 100bp DNA adder (Thermo Fisher Scientific), and run at 90V for 60 minutes in 1x TAE electrophoresis buffer. After the run was completed, DNA bands were visualized with UV light (312 nm) using a ChemiDoc Imaging System (Bio-Rad) and directly photographed.

### Tissue Pathology

At the end of the study period, EAE mice were deeply anesthetized with an IP injection of ketamine (10 mg/ml, Boehringer Ingelheim) and xylazine (1.17 mg/ml, Bayer), and transcardially perfused with saline (0.9% NaCL+ 5 M EDTA) for 5 minutes followed by 4% PFA for an additional 5 minutes. The spinal columns were isolated and post-fixed in 4% PFA at 4°C overnight. The following day, the spinal cords were extracted, washed in 1x PBS, and immersed in a 30% sucrose (in PBS) cryoprotectant solution for at least 72 hours at 4°C. Spinal cords were then embedded in optimum cutting temperature (OCT) medium in a plastic specimen holder and frozen by floating the specimen holder on 2-methylbutane (Thermo Fisher Scientific) in a petri dish on dry ice. The tissue blocks were cryo-sectioned (30 μm for *Cx3cr1*YFP^CreERT2^:R26^tdTomato^ EAE mice, 10 μm for all the other EAE experiments) using a cryostat (Leica, CM1850) with a microtome blade (Feather) onto Superfrost Plus slides (Thermo Fisher Scientific). Sections were stored at -80°C until use.

For the RFP and YFP double immunohistochemistry (IHC), sections were washed with 1x PBS + 0.3% Triton-X-100 (Sigma-Aldrich) and incubated in Bloxall (Vector Laboratories) for 10 minutes, washed 3 × 5 minutes in 1x PBS + 0.3% Triton-X-100, then blocked in 1x PBS + 0.3% Triton-X-100 + 10% normal goat serum (NGS) (Abcam) for 1 hour. Antibody incubation was done overnight at 4°C with an anti-RFP antibody (rabbit, 1:400, Abcam) in PBS + 0.3% Triton-X-100 + 10% NGS. The following day, the sections were washed 3 × 5 minutes in 1x PBS + 0.3% Triton-X-100 and incubated with biotinylated goat anti-rabbit (ready to use, Abcam) for 10 minutes, washed 3 × 5 minutes in 1x PBS + 0.3% Triton-X-100, and incubated with avidin biotin complex (ABC, Vector Laboratories) for 30 minutes, according to the manufacturer’s instruction. After washing 3 × 5 minutes in 1x PBS + 0.3% Triton-X-100, the RFP staining was visualized using brown 3,3′-diaminobenzidine (DAB) (Vector Laboratories). Sections were then washed in H_2_0 for 5 minutes, and then 1 × 5 minutes in 1x PBS. Sections were put in Bloxall for 10 minutes, washed 3 × 5 minutes in 1x PBS + 0.3% Triton-X-100, blocked with avidin and then biotin, washed 3 × 5 minutes in 1x PBS + 0.3% Triton-X-100, and then blocked with 1x PBS + 0.3% Triton-X with 10% NGS for 1 hour. Antibody incubation was done overnight at 4°C with an anti-YFP antibody (chicken, 1:1000, Abcam). The following day, sections were incubated with biotinylated goat anti-chicken (1:1000, Vector Laboratories) for 1 hour, washed 3 × 5 minutes in 1x PBS + 0.3% Triton-X-100, and incubated with ABC for 30 minutes. After 3 × 5 minutes washes in 1x PBS + 0.3% Triton-X-100, the YFP staining was visualized with blue/grey DAB (Vector Laboratories). Sections were counterstained with haematoxylin (Sigma-Aldrich) for 30 seconds and washed with dH_2_O. Sections were covered with mounting medium (Ibidi) and a cover slip placed over top (Thermo Fisher Scientific). Images were taken using an Olympus BX53 and post-processed on Fiji 2.0.0. software.

To quantify the amount of oxidative stress in EAE mice, we analysed gp91-phox expression in the spinal cord of EAE mice at 50 dpi^7^. Briefly, sections were air dried at room temperature (RT) for 25 minutes, rinsed with 1x PBS, and blocked with Bloxall for 10 minutes at RT, washed 3 × 5 minutes with 1x PBS, and incubated with the M.O.M. Immunodetection Kit (Vector Laboratories) according to the manufacturer’s instruction. Next, sections were incubated overnight at 4℃ with anti-gp91 phox antibody (mouse, 53/gp91[phox] 1:200, BD Biosciences) diluted in 1x PBS + 0.3% Triton-X with 1% NGS. The next day, the sections were washed 3 × 5 minutes in 1x PBS and incubated with a biotinylated goat anti-mouse secondary antibody (Thermo Fisher Scientific) for 1 hour at RT, washed again 3 × 5 minutes, and incubated with ABC kit (Vector Laboratories) for 1 hour at RT according to the manufacturer’s instruction. After washing 3 × 5 minutes in 1x PBS + 0.3% Triton-X, staining was visualized by applying DAB (Vector Laboratories). Sections were next washed 2x in dH_2_0 for 5 minutes. Finally, the sections were dehydrated using an alcohol gradient: 70% (2x 2 minutes), 90% (2x 2 minutes), 100% (2x 2 minutes), and xylene (2x 2 minutes), mounted (Ibidi), and coverslipped (Thermo Fisher Scientific).

To assess *in vivo* axonal loss, spinal cord sections were analysed after Bielschowsky staining, as previously performed ^8^. Briefly, sections were rehydrated in dH_2_O for 10 minutes at RT and placed in the silver nitrate solution (AgNO_3_) (10%, Sigma-Aldrich) for 20 minutes. Next, sections were washed in dH_2_O and incubated with ammonium silver solution [(AgNO_3_ (10%) with NH_3_OH (32%, Sigma-Aldrich) added till the precipitate became clear] for 20 minutes at RT. After sections turned brown, tissue was incubated in NH_3_OH solution (0.05%, Sigma-Aldrich) for 5 minutes at RT and developed in the ammonium silver solution supplemented with 100 *μ*l of developer solution [formaldehyde, 37%, Fisher Scientific), citric acid (Sigma-Aldrich), nitric acid (70%, Thermo Fisher Scientific) in dH_2_O). After 2 minutes of staining development, sections were quickly incubated in the 0.05% NH_3_OH solution for 5 minutes and washed 3 × 5 min in 1x PBS. Lastly, sections were dehydrated using an alcohol gradient: 70% (2x 2 minutes), 90% (2x 2 minutes), 100% (2x 2 minutes), and xylene (2x 2 minutes), mounted (Ibidi), and coverslipped (Thermo Fisher Scientific).

For the assessment of myeloid cell markers and axonal degeneration by immunofluorescence (IF), slides were dried and washed with dH_2_O and 1x PBS for 5 minutes each. Tissue was blocked with 1x PBS + 0.5% Triton-X-100 + 10% NGS for 1 hour at RT. For those sections stained with primary antibodies made in mouse, slides were blocked with M.O.M Mouse IgG blocking reagent (Vector Laboratories) in TBS + 0.1% Tween + 2 drops of IgG diluent for 1 hour at RT, followed by a block with M.O.M. protein diluent solution for 5 minutes at RT. The following primary antibodies were used (diluted in 1x PBS + 1% NGS and kept overnight at 4°C): anti-IBA1 (goat, 1:500, FUJIFILM Wako), anti-CX3CR1 (rabbit, 1:300, Alomone labs), anti-NDUFS4 (mouse, 1:100, Abcam or rabbit, 1:500, Novus biologicals), anti-SPP1 (goat, 1:500, R&D Systems), anti-amyloid precursor protein-APP (mouse, 1:200, Sigma-Aldrich), anti-neurofilament heavy polypeptide-NHP (rabbit, 1:500, Abcam). Of note, mouse primary antibodies were also diluted in M.O.M. protein diluent. Citric buffer (10 mM) + microwave (3 × 5 minutes) unmasking was used for the APP staining, as previously described ^9^. The next day, tissue was washed and incubated with 1:1000 secondary antibodies diluted in PBS + 0.1% Triton-X-100 + 1% NGS for 1 hour at RT. The tissue was counterstained with 1:1000 DAPI in 1x PBS for 5 minutes. Tissue was washed and embedded with mounting medium. The Leica microsystems CTR4000 confocal microscope (Leica Biosystems) was used to take confocal images of the stainings, and these were quantified using Fiji 2.0.0. software.

Quantification of *Cx3cr1*YFP^CreERT2^:R26^tdTomato^ specificity was performed on Ctrl axial spinal cord sections stained with Iba1. Quantification of the number of RFP^+^YFP^+^ (spontaneous fluorescence) cells positive or negative for Iba1 was performed on 3 spinal cord sections (4 ROIs per section) per mouse from 3 independent biological replicates. Data are expressed as percentage (%) of Iba1^+^ or Iba1^-^ cells over total number of RFP^+^YFP^+^ cells per biological replicate (±SEM).

Quantification of oxidative stress and axonal loss in EAE was obtained from analysing n ≥ 12 from spinal cord axial sections per n ≥ 4 biological replicates positive DAB area and axonal loss area respectively of equally spaced axial spinal cord sections outlined using an Olympus BX53 micro-scope with motorized stage and Neurolucida software (11.07 64-bit, Microbrightfield). Data are expressed as either percentage (%) of damaged tissue or area of positive (%) DAB staining per section (±SEM).

Quantification of axonal degeneration was performed by assessing the number of APP^+^/NHP^+^ axons on images of the posterior columns of spinal cord axial sections acquired with a 20X objective on a Leica microsystems CTR4000 confocal microscope (Leica Biosystems). Images were quantified using automatic thresholding, masking, and the image calculator tool of Fiji 2.0.0. software. Data are shown as the average number of APP^+^/NHP^+^ axons/mm^2^ (±SEM) from n=3 images per mouse (n ≥ 4 biological replicates per group).

Number of SPP1^+^ microglia (RFP^+^YFP^+^) and macrophages (RFP^-^YFP^+^), SPP1^+^ microglia (CX3CR1^+^), and SPP1^+^NDUFS4^+^ microglia (RFP^+^YFP^+^ or CX3CR1^+^) were quantified from images of the posterior columns of spinal cord axial sections acquired with a 40X objective on a Leica microsystems DMI4000 microscope (Leica Biosystems). Data are shown as the average number of positive cells/mm^2^ (±SEM) from n=5 images per mouse (n ≥ 2 biological replicates per group).

### Suspension Mass Cytometry (CyTOF)

At 50 dpi, the first group of EAE NDUFS4 KO mice (n=4) for CyTOF were deeply anaesthetized with IP injections of ketamine (10 mg/ml, Boehringer Ingelheim) and xylazine (1.17 mg/ml, Bayer) and then transcardially perfused with ice-cold 1x PBS + 0.1% EDTA. Spinal cords were extracted from the spinal columns using a 5 mL syringe filled with ice-cold sorting buffer [DMEM no phenol red (Thermo Fisher Scientific) + 25mM HEPES (Sigma-Aldrich) + 5% dialyzed FBS (Thermo Fisher Scientific) + 1mM EDTA (Honeywell Fluka) + 1x glutaMAX (Thermo Fisher Scientific)]. Spinal cords were mechanically dissociated in a glass Dounce tissue homogenizer with 7 mL of cold homogenization buffer (sorting buffer + 10 mg/mL of DNAse). After tissue dissociation, the suspension was filtered through a pre-wet 40 μM strainer and the homogenizer rinsed with 2 mL of homogenization buffer. Samples were then transferred to 15 mL Falcon tubes and 2.7 ml of 90% Percoll (GE Healthcare) in 10x PBS (Thermo Fisher Scientific) was added to each sample to remove myelin and debris. The 15 mL Falcon tubes were then inverted 10 times gently and the samples were centrifuged (CENTRIFUGE MODEL) at 800 × *g* for 20 minutes at 4°C with a brake speed of 0. Myelin debris visibly layered at the surface was carefully removed with a pipette. Ice-cold buffer [95% autoMACS rinsing solution (Miltenyi Biotec) and 5% MACS BSA (Miltenyi Biotec)] was added to the samples (to fill the 15 mL tubes) and samples were centrifuged (CENTRIFUGE MODEL) at 800 × *g* for 5 minutes at 4°C to remove the remaining Percoll. Pellets were resuspended in sorting buffer (1 mL+7 mL) and then further centrifuged at 800 × *g* for 5 minutes at 4°C.

The cell pellet was resuspended in 100 μL of cold buffer and 10 μL of CD45 positive selection MicroBeads (Miltenyi Biotec) were added to each sample and mixed well. The samples were incubated at 20 minutes on ice and gently mixed every 5 minutes. After incubation, 2 mL of cold buffer was added to wash the samples and then centrifuged at 300 × *g* for 10 minutes at 4°C. After centrifugation, the supernatant was removed with a pipette, and the pellet was resuspended in 500 μL of cold buffer. The LS MACS columns (Miltenyi Biotec) were placed in the QuadroMACS Separator (Miltenyi Biotec) on the MACS MultiStand (Miltenyi Biotec) and rinsed 3 × 3 mL of cold buffer. After the columns were rinsed, the 500 μL cell suspension was added onto the column. Unlabelled (i.e., CD45^-^) cells from the filtrate were collected and the column was washed 3 × 3 mL of cold buffer. Finally, 5 mL of cold buffer was added to the column and subsequently flushed with a plunger to isolate the CD45^+^ cells. The samples were centrifuged at 300 × *g* for 5 minutes at 4°C. The supernatant was removed, and the cells were resuspended in 50 μL of Maxpar PBS (Fluidigm). A working solution (1:500) of 5 μM Cell-ID Cisplatin (Fluidigm) was prepared and 50 μL was added to the cells (1:1 mix) for a final concentration of 2.5 μM. The cells were incubated for 90s and the cisplatin was quenched with 5 mL of Maxpar cell staining buffer (Fluidigm). The samples were centrifuged at 300 × *g* for 5 minutes at 4°C, the supernatant removed, and the cells re-suspended in 1 mL of 1x Fix I buffer (Fluidigm) diluted in Maxpar PBS and left to incubate for 10 minutes at RT. The samples were centrifuged at 1000 × *g* for 7 minutes at RT, the supernatant was removed, and the cells were resuspended in 1 mL of 1x Barcode perm buffer (Fluidigm) then re-centrifuged at 1000 × *g* for 7 minutes at RT. The supernatant was removed, and the cells were resuspended in 800 μl of 1x Barcode perm buffer. Then, 100 μL of a Cell-ID palladium (Pd) unique barcode (Fluidigm) was added to each sample and left to incubate for 30 minutes at RT. Samples were gently mixed after 15 minutes. Following barcoding, the samples were centrifuged at 1000 × *g* for 7 minutes at RT, the supernatant was removed, and the cells were resuspended in 2mL of Maxpar cell staining buffer. The samples were re-centrifuged at 1000 × *g* for 7 minutes at RT, the supernatant was removed, and the cells were resuspended in 100 μL of Maxpar cell staining buffer. All the barcoded samples were combined into one single 5 mL polypropylene tube (BD Biosciences). Each samples tube was washed with 500 μL of Maxpar cell staining buffer, transferred to the single tube (containing the barcoded samples), and centrifuged at 1000 × *g* for 7 minutes at RT. The supernatant was removed, the cells were resuspended in 50 μL of Maxpar cell staining buffer and CD16/CD32 Fc receptor block (BD Biosciences) at 1:25 was added. The sample was left to incubate for 10 minutes at RT. After 10 minutes, 50 μL of the surface antibody cocktail (Supplementary Data 6) was added to the sample. The sample was mixed with gentle pipetting and incubated for 15 minutes at RT. At 15 minutes, the sample was gently mixed and left to incubate for an additional 15 minutes. After surface antibody incubation, 2 mL of Maxpar cell staining buffer was added to the sample and it was centrifuged at 1000 × *g* for 7 minutes at RT. The supernatant was removed, 2 mL of Maxpar cell staining buffer was added to the sample, and it was re-centrifuged at 1000 × *g* for 7 minutes at RT. The supernatant was removed, and the cells were fixed in 1 mL of 1x Maxpar Fix I buffer for 15 minutes at RT. Then, 2 mL of Maxpar Perm-S buffer (Fluidigm) was added and the sample was centrifuged at 1000 × *g* for 7 minutes at RT. The supernatant was removed, 2 mL of Maxpar Perm-S buffer was added and the sample was re-centrifuged at 1000 × *g* for 7 minutes at RT. The supernatant was removed (leaving behind ∼50 μL) and 50 μL of the cytoplasmic/secreted protein antibody cocktail (Supplementary Data 6) was added. The sample was gently mixed with a pipette and left to incubate for 30 minutes at room temperature (at 15 minutes the sample was gently mixed). After incubation, the cells were washed with 2 mL of Maxpar cell staining buffer, and centrifuged at 1000 × *g* for 7 minutes at RT. The supernatant was removed, 2 mL of Maxpar cell staining buffer was added to the sample, and it was re-centrifuged at 1000 × *g* for 7 minutes at RT. The supernatant was removed, and the cells were fixed in 1 mL of 1.6% PFA for 10 minutes at RT. After 10 minutes, the cells were centrifuged at 1000 × *g* for 7 minutes at RT. The supernatant was removed, and the cells were incubated with 1 mL of Cell-ID Intercalator-IR diluted in Maxpar Fix and Perm buffer [1:1000 dilution of 125 μM stock solution] (Fluidigm) for 1 hour at RT. After incubation, the cells were centrifuged at 1000 × *g* for 7 minutes at RT. The supernatant was removed, 2 mL of Maxpar cell staining buffer was added to the sample, and it was re-centrifuged at 1000 × *g* for 7 minutes at RT. The supernatant was removed, 1 mL of Maxpar cell staining buffer was added to the sample, and it was re-centrifuged at 1000 × *g* for 10 minutes at RT. At this stage, the supernatant was removed leaving between 50-100 μL of solution covering the cell pellet. The sample was stored for 48 hours at 4°C until acquisition. The following day the other group of EAE NDUFS4 KO mice (n=4) were processed for CyTOF following the same protocol. This set of barcoded samples was stored for 24 hours at 4°C until acquisition. The day of acquisition, the samples were combined into a single tube of 8 barcoded samples. The cells were washed in 1 mL of Maxpar Cell Acquisition Solution (Fluidigm) and centrifuged at 1000 × *g* for 7 minutes at RT. The supernatant was removed, and the samples were left on ice until acquisition. Directly prior to acquisition, the cells were resuspended in 1 mL of 0.1X EQ Four Element Calibration Beads (Fluidigm) diluted in Maxpar Cell Acquisition Solution, passed through a 40 μM filter, and then analysed on a Helios™ mass cytometer using a WB injector. The data files were debarcoded using Fluidigm software (v7.0.8493) with a Mahalanobis distance of 2.50 then concatenated and exported as FCS files. FCS files were imported into FlowJo (BD Biosciences), and live cells were selected using Boolean gating. Then, each sample was gated for myeloid cells (CD45^+^CD11b^+^) and lymphoid cells (CD45^+^CD11b^-^), exported as individual FCS files, and further analysed in R version 4.1.3.

For the EAE mice that received the small molecule treatments (n=3 mice/treatment group), at 30 dpi the first group of mice (n=6) for CyTOF were deeply anaesthetized with IP injections of ketamine (10 mg/mL, Boehringer Ingelheim) and xylazine (1.17 mg/ml, Bayer) and then transcardially perfused with ice-cold 1x PBS + 0.1% EDTA. Then, the spinal cords were processed for CyTOF as described for the NDUFS4 KO EAE mice. The first set of samples (n=6) were stored for 48 hours at 4°C until acquisition. The other set of samples (n=6) were processed for CyTOF the following day and stored for 24 hours at 4°C until acquisition. The data files were debarcoded using Fluidigm software (v7.0.8493) with a Mahalanobis distance of 8.0 then concatenated and exported as FCS files. FCS files were imported into FlowJo and live cells were selected using Boolean gating. Then, each sample was gated for myeloid cells (CD45^+^CD11b^+^) and lymphoid cells (CD45^+^CD11b^-^), exported as individual FCS files, and further analysed in R version 4.1.3.

All packages are publicly available at the Comprehensive R Archive Network (https://cran.r-project.org), the Bioconductor project (http://bioconductor.org), or their respective GitHub pages. The original code is available at https://github.com/regan-hamel/ComplexI.

## Supplementary Data

**Supplementary Data 1.** ScRNAseq of myeloid cells isolated from *Cx3cr1*YFP^CreERT2^:R26^tdTomato^ EAE mice. Tabs show cell type variation, differentially expressed genes (DEGs), and GO terms for each cluster.

**Supplementary Data 2.** ScRNAseq of specific myeloid cells sub-clusters isolated from *Cx3cr1*YFP^CreERT2^:R26^tdTomato^ EAE mice. Tabs show differentially expressed genes (DEGs) and GO terms for subclusters 4.1-3 and 6.1-5, as well as DEGs for MAMS.

**Supplementary Data 3.** LC-MS analysis of the intracellular metabolome of myeloid cells isolated from *Cx3cr1*YFP^CreERT2^:R26^tdTomato^ EAE mice. Tabs report original raw data, normalized data and data depicted in the Volcano plots.

**Supplementary Data 4**. Multivariate analysis of the LD-REIMS metabolomics data of the EAE mice. Tabs show the average normalized intensity values of metabolites of interest driving the stratification of various regions-of-interest. Species names have also been assigned alongside the corresponding m/z values where appropriate.

**Supplementary Data 5**. ScRNAseq of CNS cells isolated from WT and *Nd6* EAE mice. Tabs show differentially expressed genes (DEGs) and GO terms for each cluster.

**Supplementary Data 6**. CyTOF panel and antibodies.

